# Human brain organoids record the passage of time over multiple years in culture

**DOI:** 10.1101/2025.10.01.679721

**Authors:** Irene Faravelli, Noelia Antón-Bolaños, Anqi Wei, Tyler Faits, Abhishek Sampath Kumar, Sophia Andreadis, Rahel Kastli, Marta Montero Crespo, Mara Steiger, Daniel Leible, Elizabeth Zhang, Bobae An, Yaron Meirovitch, Sayara Silwal, Sung Min Yang, Alexander Kovacsovics, Xian Adiconis, Helene Kretzmer, Joshua Z. Levin, Edward S. Boyden, Jeff Lichtman, Aviv Regev, Alexander Meissner, Paola Arlotta

**Author notes:** These authors contributed equally (Irene Faravelli, Noelia Antón-Bolaños). These authors contributed equally (Anqi Wei, Tyler Faits).

## Abstract

The human brain develops and matures over an exceptionally prolonged period of time that spans nearly two decades of life. Processes that govern species-specific aspects of human postnatal brain development are difficult to study in animal models. While human brain organoids offer a promising *in vitro* model, they have thus far been shown to largely mimic early stages of brain development. Here, we developed human brain organoids for an unprecedented 5 years in culture, optimizing growth conditions able to extend excitatory neuron viability beyond previously-known limits. Using module scores of maturation-associated genes derived from a time course of endogenous human brain maturation, we show that brain organoids transcriptionally age with cell type-specificity through these many years in culture. Whole-genome methylation profiling reveals that the predicted epigenomic age of organoids sampled between 3 months and 5 years correlates precisely with time spent *in vitro,* and parallels epigenomic aging *in vivo*. Notably, we show that in chimeric organoids generated by mixing neural progenitors derived from “old” organoids with progenitors from “young” organoids, old progenitors rapidly produce late neuronal fates, skipping the production of earlier neuronal progeny that are instead produced by their young counterparts in the same co-cultures. The data indicate that human brain organoids can mature and record the passage of time over many years in culture. Progenitors that age in organoids retain a memory of the time spent in culture reflected in their ability to execute age-appropriate, late developmental programs.

## INTRODUCTION

Human brain development is orchestrated by transcriptional programs defined by spatially and temporally regulated waves of gene expression. These gene expression patterns are tightly coordinated by dynamic changes in the activity of regulatory elements and epigenomic remodeling, ensuring the timely emergence of distinct neural cell types and the progressive maturation of the nervous system. In humans, brain development and maturation proceed at a markedly slower pace than in most other species, with cortical neurons requiring years to reach full maturity. This protracted timeline is preserved in all cell types generated *in vitro* from human pluripotent stem cells (hPSCs), including cortical neurons, pointing at a cell-intrinsic clock that sets the pace of brain development, although its molecular mechanisms and functional significance remain to be elucidated(Ciceri et al., 2024; Del Dosso et al., 2020). Recent work has shown the importance of epigenetic barriers that enforce the slow timing of human neuronal maturation(Ciceri et al., 2024), and the role of species-specific rates of mitochondrial metabolism in influencing developmental tempo(Casimir et al., 2024). Human organoids could in principle be used to study these processes of human brain maturation, but existing models largely recapitulate earlier developmental processes, and most studies do not follow organoids over a sufficiently long timespan. One report cultured organoids for up to 694 days, and used bulk RNA-sequencing and methylation arrays to profile whole organoids(Gordon et al., 2021). However, without profiling of the individual cell types present within these systems, and absent longer timelines of maturation, we currently lack understanding of the mechanisms by which the many cell types of the human brain measure and record time.

In this study, we developed human cortical organoids for over 5 years in culture and integrated single-cell transcriptional information with epigenetic, structural and functional data to build a comprehensive map of development and maturation across an unprecedented time span in culture. The data indicate that human brain organoids record the passage of time following endogenous milestones and using similar epigenetic mechanisms. Importantly, progenitors in old organoids can recall the passage of time, like a memory of development that has already happened, to generate late progeny without repeating early developmental steps. The work demonstrates that cells in organoids can develop over unprecedented time periods and are capable of both recording and recalling their developmental time. These systems provide a wealth of data on largely inscrutable periods of postnatal development of the human brain.

## RESULTS

### Brain cortical organoids age appropriately in culture

Development and maturation of human brain organoids follows the notoriously slow, neotenous pace of the endogenous human brain, which requires many years to reach adulthood(Fenlon, 2021; He et al., 2023; Uzquiano et al., 2022; Velasco et al., 2019). Thus far, organoids have mostly been studied over timespans of months, and it remains unclear whether they can continue to mature and age in culture over many years to mimic aspects of postnatal maturation of the human brain that otherwise remain largely experimentally inaccessible. To investigate this possibility, we have cultured human organoids of the cerebral cortex for over 5 years. To begin to characterize the cellular and molecular features of these long-term cultures, we used single-cell RNA sequencing (scRNA-seq) to profile 26 single organoids collected at 6 months, 9 months, 1 year, 1.5 years, 2 years, 3 years and 4 years, and integrated this data with a dataset from 51 single organoids that we had previously profiled at multiple timepoints from 3 months through 6 months(Antón-Bolaños et al., 2024; Uzquiano et al., 2022; Velasco et al., 2019) (see **Methods**; **Fig. 1A-B**, n= 77 replicates in total from 26 new datasets + 51 previously generated, n= 257,044 cells).

**Figure 1.**
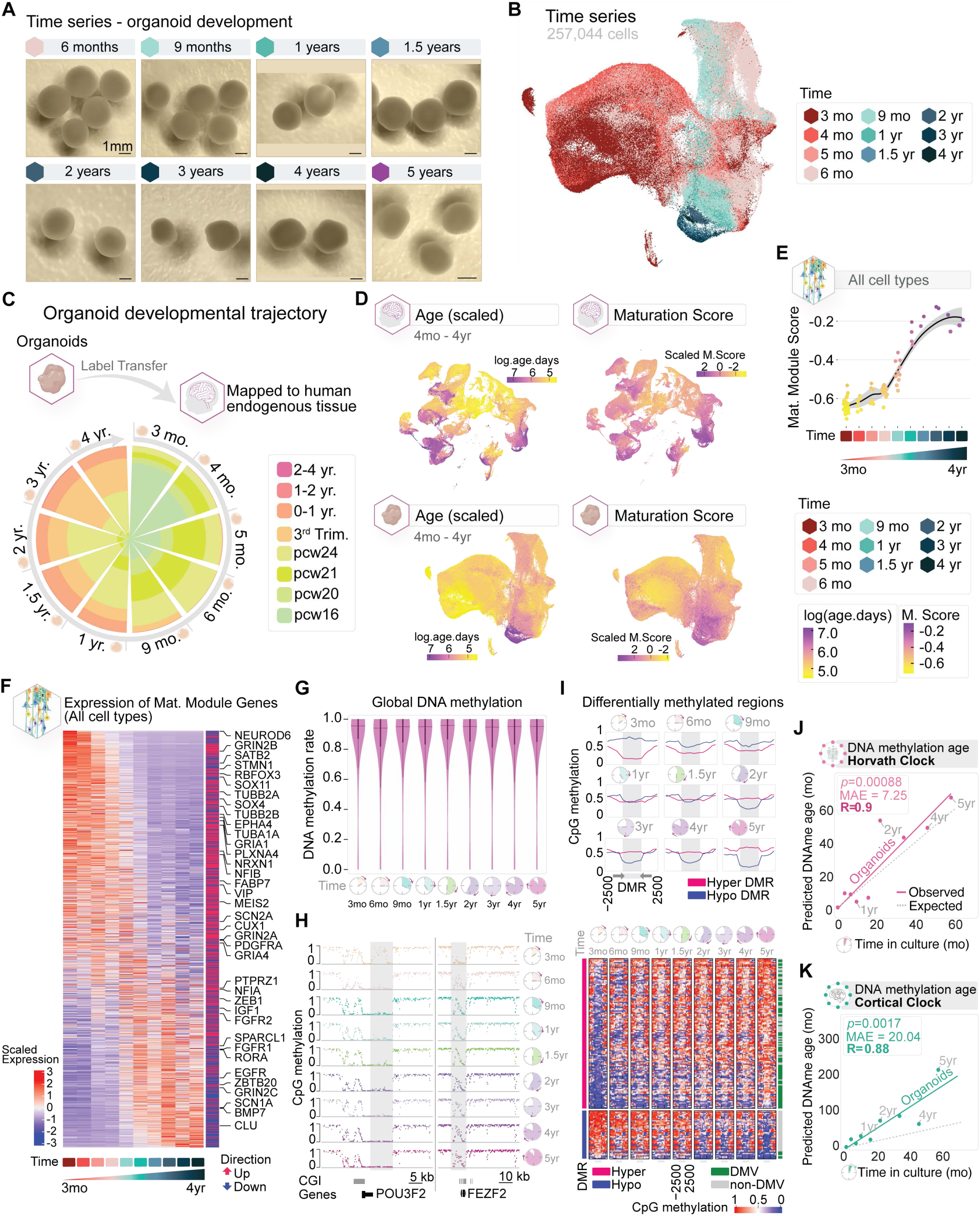
**A**, Brightfield representative images of the organoid time-series. Scale bar: 1 mm. **B**, UMAP plot showing the distribution of cortical organoid ages. (n=257,044 cells across 10 timepoints.). **C**, Rose plot showing the age range of the endogenous perinatal human tissue(Velmeshev et al., 2023) that cells from each cortical organoid timepoint mapped to via label transfer. Organoid cells that mapped to fetal 2nd trimester were mapped to a second endogenous fetal dataset(Trevino et al., 2021) to add additional granularity to their mapped age. The initial tissue reference dataset spanned from fetal 2nd trimester to 2-4 years postnatal age, while the follow-up reference spanned from post-conception week (PCW) 16-PCW 24. **D**, UMAP plots showing (subsetted and renormalized) data from endogenous perinatal tissue (Velmeshev et al., 2023), and cortical organoids. Plots are colored by the log-scaled age of each sample/organoid and the Z-scored Maturation Score of each cell. Maturation Scores represent the overall expression of the genes belonging to a Multicellular Program (MCP) – MCP4 – identified by DIALOGUE (see **Methods** for details). **E**, Scatterplots showing the mean Maturation Score (y-axis) of all cells within each organoid, grouped by timepoint (x-axis). Smoothed conditional means (black lines) with default 95% confidence intervals (grey regions) were calculated by the geom_smooth() function in R. Points were given slight x-axis jitter to improve visibility; horizontal position within a timepoint does not indicate differences in age. **F**. Heatmap showing the expression of all genes contributing to MCP4 in organoids across time. Values for each timepoint represent the mean of the counts-per-million of the pseudobulked count matrices for each organoid within that timepoint, Z-scored for each gene. Genes are ordered by the weighted expression over time; genes expressed mostly at earlier timepoints are at the top, genes expressed mostly at later timepoints are at the bottom. The color bar on the right indicates whether the gene was marked as in the “up” (red) or “down” (blue) aspect of MCP4. Labeled genes were manually selected to show important biological features over time. **G**, Global CpG DNA methylation levels. The line in the violin plot indicates the median of the distribution. **H**, Genome browser tracks showing methylation profiles of DMRs at the POU3F2 and FEZF2 loci across all time points. DMR regions are highlighted (grey boxes). CGIs, and genes are annotated below. **I**, Average CpG methylation profiles across n=164 hyper- and hypo-DMRs identified between 3-month and 5-year organoids that show strong correlation with time-in-culture (Pearson correlation coefficient > 0.6). Heatmap showing methylation profiles of all time points across n=164 hyper- and hypo-DMRs strongly correlated with time-in-culture. Overlaps of DMRs with DMVs are indicated on the right. The dashed line denotes the DMR, and the regions to its right and left are +/- 2.5kb. **J**-**K**, The predicted DNA methylation age for the set of Horvath (**J**) or Cortical clock (**K**) CpGs (approximately 350) is presented. The x-axis represents the chronological age of the organoids (in months), while the y-axis shows the predicted DNA methylation age in the organoids (also in months). The dashed line indicates perfect correlation, while the solid line represents linear regression with the R value. MAE indicates the median absolute error.

To correlate transcriptional changes in organoids over years *in vitro* with those occurring in the human brain over years *in vivo*, we leveraged existing scRNA-seq datasets of the endogenous human cortex spanning developmental timepoints from the second trimester(Trevino et al., 2021) to postnatal stages of development, up to 4 years(Velmeshev et al., 2023). Using the endogenous datasets as reference, we used label transfer to estimate the transcriptional “age” of organoid cells (see **Methods**). Analyzing each organoid timepoint separately, reference-based label transfer revealed that early organoids (3- to 6-month) predominantly mapped to the second trimester human fetal brains, while later stages (9-month to many years) progressively shifted towards mapping to late prenatal and postnatal stages (**Fig. 1C**). This suggested a temporal alignment with in *vivo* developmental trajectories, with postnatal-like signatures emerging in organoids after 9 months in culture.

To precisely assess transcriptional maturation, we first identified gene modules correlated with age in the endogenous human cortex. We applied DIALOGUE(Jerby-Arnon and Regev, 2022), a computational framework for deriving multicellular programs (MCPs) of co-regulated genes across different cell types, to an *in vivo* human brain cortex single-cell dataset spanning from the second trimester up to 4 years of age (derived from(Velmeshev et al., 2023), **Fig. 1D-F**). We then projected these same maturation modules onto the organoid time-series dataset ranging from 3 months to 4 years in culture. This analysis revealed a strong correlation between *in vivo* age-related module scores and duration of *in vitro* development, showing that organoids display the same coordinated, age-linked transcriptional programs observed *in vivo*. The findings show that cells in organoids preserve a sequential progression of gene expression programs associated with progressive maturation *in vivo*.

To define whether organoids show appropriate age-related epigenetic programs over the years, we next performed whole-genome bisulfite sequencing (WGBS) of single organoids at different ages. DNA methylation, particularly at age-associated CpG sites, serves as a powerful molecular readout of biological age(Moqri et al., 2025). We sampled organoids at nine time points from 3 months to 5 years in culture (**Fig. 1G-F, Supplementary Fig. 1A**). *In vivo*, across multiple tissues, the methylome is dynamic during early development but then reaches stable high levels of DNA methylation that are maintained over time. In contrast, in 2D culture systems, the methylome is more unstable and epigenomic drift is common(Cruickshanks et al., 2013; Edgar et al., 2022; Franzen et al., 2021; Nishino et al., 2011; Zhou et al., 2018a). WGBS profiling of organoids over our extended time-course showed that organoids maintain global stability of DNA methylation patterns (**Fig. 1G**), similar to the high and stable methylation levels observed in postnatal and adult cortical tissue. Pairwise comparisons of genome-wide methylation revealed a strong correlation between adjacent time points (Pearson r = 0.89–0.94), supporting the preservation of the methylation landscape in multi-year cultures (**Supplementary Fig. 1B**). Global CpG methylation remained high across all time points; both partially methylated domains (PMDs), predominantly affected by methylation drift *in vitro*, and highly methylated domains (HMDs) retained near-complete methylation at all ages across the 5 years (**Supplementary Fig. 1C**). DNA methylation valleys (DMVs), which often demarcate developmental regulatory regions, remained stable (**Supplementary Fig. 1D**). CpG methylation at repetitive elements similarly remained stable over time, further underscoring the global preservation of the methylome in long-term organoid cultures (**Supplementary Fig. 1E**). Together, these findings indicate that the longitudinal methylation landscape in organoids during extended culture resembles epigenetic features of endogenous tissues.

Despite the overall stability of methylation in endogenous tissues, regulation of methylation state at specific developmentally-relevant genes are required for cell fate specification and overall development. To determine whether developmentally-regulated loci undergo regulated changes over time in organoids, we examined CpG methylation at key cortical genes involved in fate acquisition and layer specification—*FEZF2* and *TBR1* for deep-layer projection neurons, *POU3F2* for upper-layer neuron identity, and *NEUROD2* for broader excitatory neuron differentiation. Each of these loci showed progressive, time-dependent methylation gains, particularly within CpG islands and DMVs (**Fig. 1H** and **Supplementary Fig. 1H**).

To examine temporal changes over organoid development, we used differential methylation analysis between the two endpoints, 3-month and 5-years, to identify differentially methylated regions (DMRs, **Supplementary Fig. 1F**). DMRs highlight parts of the genome where methylation actively changes between conditions; in this case, between early and late cultures. We found 213 age-correlated DMRs, including both hyper-DMRs, which gained methylation over this period, and hypo-DMRs, which lost methylation (**Supplementary Fig. 1G**). We then examined the behavior of these DMRs over the intermediate timepoints, and found that rather than arising abruptly or stochastically, these methylation shifts emerged continuously over the culture period, suggesting a tightly regulated epigenetic program (**Fig. 1I**). The relatively small number of DMRs and their gradual emergence at intermediate time points suggest coordinated regulation that is non-fluctuating rather than random drift. Functional annotation revealed that DMRs were enriched in epigenetic features of known developmental importance, including CpG islands, DNA methylation valleys, and gene regulatory elements. Notably, the genes associated with these DMRs include many known to play key roles in the developing neocortex, such as P*AX6, HES6, EMX1, EMX2, NES, TBR1, EOMES, SATB2, POU3F2* (**Fig. 1I**). While it is difficult to quantify the exact number of methylation changes across comparable *in vivo* developmental windows due to the limited *in vivo* data available in the literature, the number and nature of DMRs observed in organoids, and the fact that they are enriched in developmentally relevant loci and show coordinated progression, are consistent with regulated *in vivo* methylation programs rather than stochastic methylation drift.

Finally, we examined CpA methylation (mCA), a non-CpG modification that is a hallmark of neuronal maturity(Luo et al., 2016). In neurons, mCA accumulates postnatally and is believed to contribute to long-term gene silencing and transcriptional fine-tuning, making it a valuable molecular marker of neuronal maturation(Luo et al., 2016). Global mCA levels increased steadily in organoids over the first year (**Supplementary Fig. 1I**) and remained stable thereafter. Importantly, when we examined a set of 1383 human fetal forebrain super-enhancers that gain mCA during postnatal neuron development *in vivo*(Luo et al., 2016), we found progressive mCA at these sites over time in organoids. This indicates that neuron-specific, non-CpG methylation pathways are not only active in our *in vitro* system, but are temporally regulated at appropriate target loci (**Supplementary Fig. 1G, K)**. Together, these results imply that the ongoing methylation dynamics observed in organoids over extended culture may reflect *in vitro* recapitulation of endogenous development and maturation epigenetic programs.

Prior work has identified CpG sites that show reproducible gains or losses in methylation over time(Christensen et al., 2009). These sites have been used to create age clocks, capable of correlating methylation at these informative sites with the temporal age of the tissue, *in vivo*(Horvath, 2013). To evaluate whether organoids capture features of biological aging, we applied two epigenetic clocks: the Horvath pan-tissue clock(Horvath, 2013) and a human cortex-specific DNA methylation clock(Shireby et al., 2020) (**Fig. 1J-K**). In both models, predicted DNA methylation age (DNAm age) tracked closely with culture time (R = 0.88–0.90), with median absolute errors of 7.25 months and 20.04 months, respectively (**Fig. 1J-K**). This indicates that organoids recapitulate time-resolved epigenetic aging signatures of endogenous tissues. To verify that the observed methylation changes were not primarily driven by proliferation, we examined methylation at solo-WCGW tetranucleotide sites, which correlates with mitotic division history, and confirmed that methylation at these sites remained stable (**Supplementary Fig. 1L**).

Collectively, these findings demonstrate that organoids are capable of preserving robust, time-resolved transcriptional and DNA methylation programs that are associated with tissue aging *in vivo*. The capacity to recapitulate transcriptional and epigenetic signatures of *in vivo* maturation across 5 years *in vitro* establishes organoids as a powerful model system to study the biology of both the prenatal and postnatal human brain.

### Activity-permissive culture conditions enable long-term maintenance of excitatory neurons with enhanced maturation

It is well known that not all cell types survive equally well in organoids. Using scRNA-seq data collected from single organoids over time, we found that although all cell types were generated in cortical organoids (**Supplementary Fig. 2**), and astrocytes and different types of progenitors remained well represented for 4 years in culture, all neuronal populations progressively declined in proportion over time. It is possible that the neuronal-specific decline observed by scRNA-seq be due to technical limitations associated with neuronal fragility and death upon tissue dissociation; however, this loss is consistent with previous findings that neurons decline over time in culture, likely due to the lack of spontaneous activity, which conventional media does not support(Bardy et al., 2015).

Given the critical role of neuronal activity for neuronal maturation, function and survival, we modified a previously established medium designed to preserve spontaneous activity (BrainPhys(Bardy et al., 2015)), to test whether it could support neuronal survival in organoids past 6 months. We tested a modified BrainPhys-based medium, which we termed Activity Permissive Medium (APM; see **Methods**), and cultured organoids in APM starting from DIV 70 (**Fig. 2A**). We found that, at 6 months, neurons from organoids cultured in APM showed a marked increase in gene module scores for synapse-related genes (see **Methods**) (**Fig. 2B, Supplementary Fig. 3A-D**). Immunohistochemistry confirmed expression of c-FOS, an early marker of neuronal activity, in callosal projection neurons at 9 months (**Fig. 2C**). Gene ontology analysis further supported enrichment of GO terms related to synapse organization and function (**Supplementary Fig. 3E**). Together, these data indicate that APM-cultured organoids show transcriptional responses consistent with the presence of neurons displaying increased activity.

**Figure 2.**
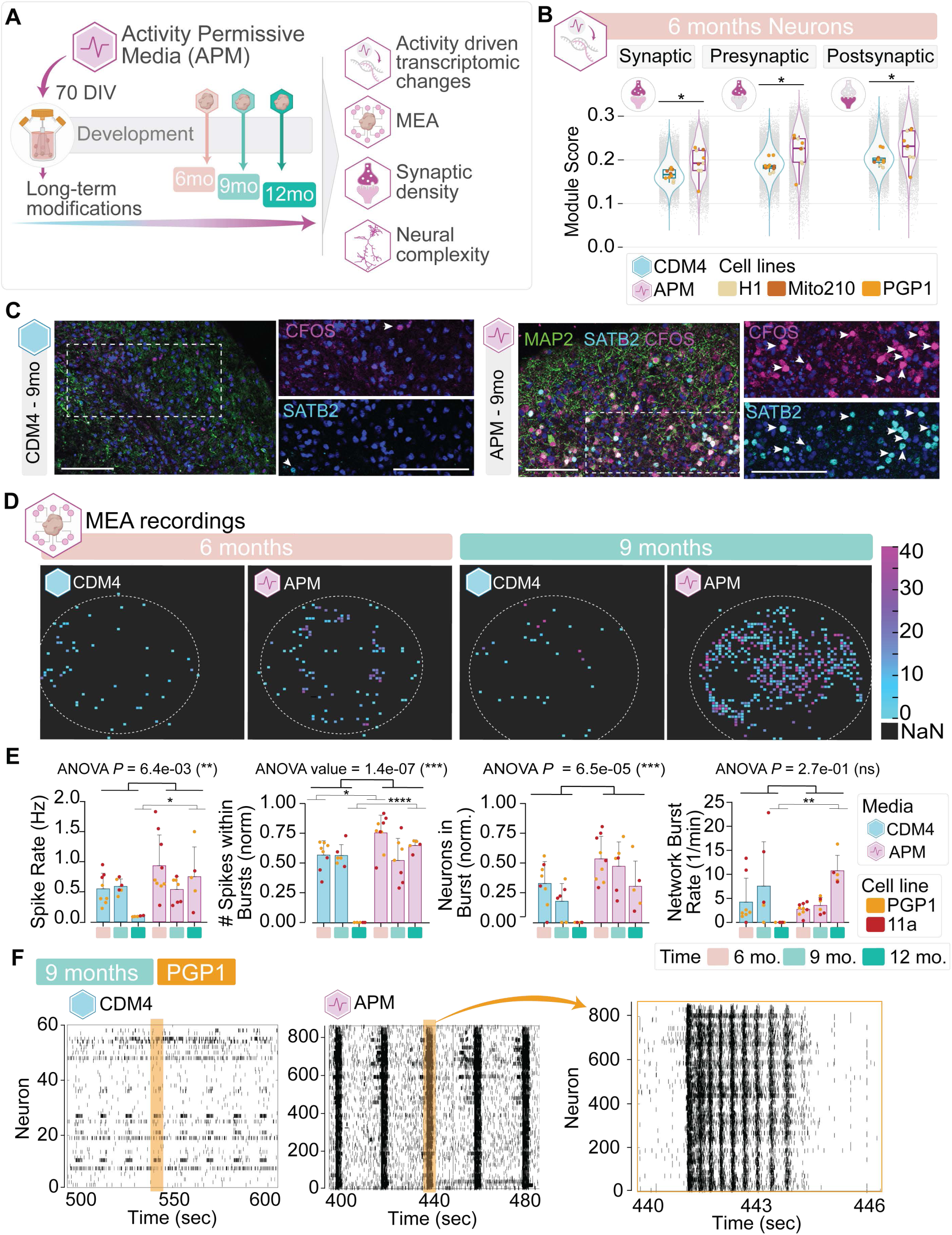
**A**, Schematics. **B**. Expression of three SynGO synaptic gene sets is higher in neurons of APM-treated organoids at 6 months (CDM4: 29,987 cells from n = 10 organoids; APM: 22,422 cells from n = 9 organoids). Violin plots show the module score distribution for synapse (GO:0045202), presynapse (GO:0098793), and postsynapse (GO:0098794) gene sets retrieved from the SynGO knowledge base across neuronal populations annotated at 6 months. Averaged module scores for individual organoids are shown as points, colored by genetic backgrounds. The bottom and top edges of the boxes represent the 25th and 75th percentiles, with the middle line representing the median and whiskers spreading the 1.5 x interquartile range (IQR) from the hinges. Module scores were calculated by using Seurat’s AddModuleScore function. Linear mixed-effects models were used to model the scores by including which organoid the cell belongs to and the genetic background of that organoid as random effects. A weight factor was incorporated to adjust for cell count differences in the neuronal partition across organoids. Models with or without treatment as an additive fixed effect were evaluated by anova function. FDR-adjusted *P* values of 2.22 × 10^-2^ (synapse), 2.22 × 10^-2^ (presynapse), and 2.58 × 10^-2^ (postsynapse) are indicated by asterisk as follows: *P* < 0.05 (*); *P* < 0.01 (**); *P* < 0.001 (***); *P* < 0.0001 (****). **C**, Immunohistochemistry of CDM4 and APM organoids at 9 months showing cFOS (early activity marker) and SATB2 (callosal projection neuron marker). Scale bar: 100 μm. **D**, Representative images of multielectrode array (MEA) recordings. **E**. For each measured metric by MEA, a Type III two-way ANOVA test was performed to assess the effects of Treatment, Age, and their interaction Treatment:Age. For cases where quantifications of more than one cell line were available, the genetic background was included as a fixed effect. ANOVA *P* value is shown to denote the significance of the Treatment effect on the measured metric, as represented by asterisks. Post-hoc Tukey tests were performed to determine at which age the treatment significantly differ and adjusted *P* values by Tukey’s HSD method were represented by asterisks. **F**. Representative raster plots of 9-month CDM4 and APM organoids.

To directly assess the effect of APM exposure on circuit activity, we recorded spontaneous extracellular activity using a 3D multielectrode array (MEA) platform (see **Methods**, previously described(Antón-Bolaños et al., 2024)) at 6, 9 and 12 months of differentiation (total of 19 single organoids derived from two different donors, **Fig. 2D**). We compared organoids that were cultured continuously in APM from 70 DIV to control organoids derived from the same lines that were cultured in conventional media (CDM4) and transferred to APM two weeks before the recordings (as previously described(Paulsen et al., 2022)). For each organoid, we recorded a minimum of 20 minutes of spontaneous activity and identified spikes post hoc using the kilosort algorithm(Pachitariu et al., 2024), which we optimized for application to organoids (see **Methods**).

The majority of recorded organoids in both APM and CDM4 conditions exhibited periodic bursts of highly correlated activity, known as “network bursts,”. We verified that this bursting behavior could be abolished by the application of the NMDA and AMPA glutamate receptor blockers, D-AP5 and DNQX, suggesting that the bursts are mediated by synaptic glutamatergic transmission. Recorded activity was abolished by the application of TTX, indicating that the observed signals were mediated by action potentials (**Supplementary Fig. 4A, B**). However, we found significant differences in neuronal activity between CDM4 and APM culture conditions across all three differentiation stages. Across all ages, APM organoids displayed a higher spike rate and increased network burst frequency (**Fig. 2E, Supplementary Fig. 4C, D**); additionally, APM organoids had more spikes per burst, and a greater number of neurons participated in each burst (**Fig. 2E, Supplementary Fig. 4C, D**). The most pronounced differences between the two conditions were observed at the 12-month mark: organoids cultured in CDM4 for 12 months consistently failed to exhibit any network bursts (5 out of 5), whereas all APM organoids at 12 months exhibited robust network bursts (5 out of 5) (**Fig. 2E, Supplementary Fig. 4C, D**). Notably, starting from 9 months of differentiation, APM organoids displayed more complex bursting patterns, while CDM4 organoids showed increasingly sparse activity over the same period (**Fig. 2E, F** and **Supplementary Fig. 4C, D**). Overall, our results indicate that neuronal circuits in APM-cultured organoids develop and sustain enhanced activity for at least one year in culture.

To investigate whether the observed increase in activity was accompanied by changes in ultrastructural features, we conducted quantitative electron microscopy (EM) analysis on organoids cultured in either CDM4 or APM at 6 and 12 months (n=3 organoids per time point and condition; n=2 genetic backgrounds at 6 months: H1 and 11a, and n=3 genetic backgrounds at 12 months: H1, 11a, and PGP1, for both CDM4 and APM conditions). We acquired large 2D panoramic images (approximately 350 µm x450 µm) that covered a representative portion of each organoid, spanning from the surface to the core, to examine structures at different depths (**Fig. 3A**).

**Figure 3.**
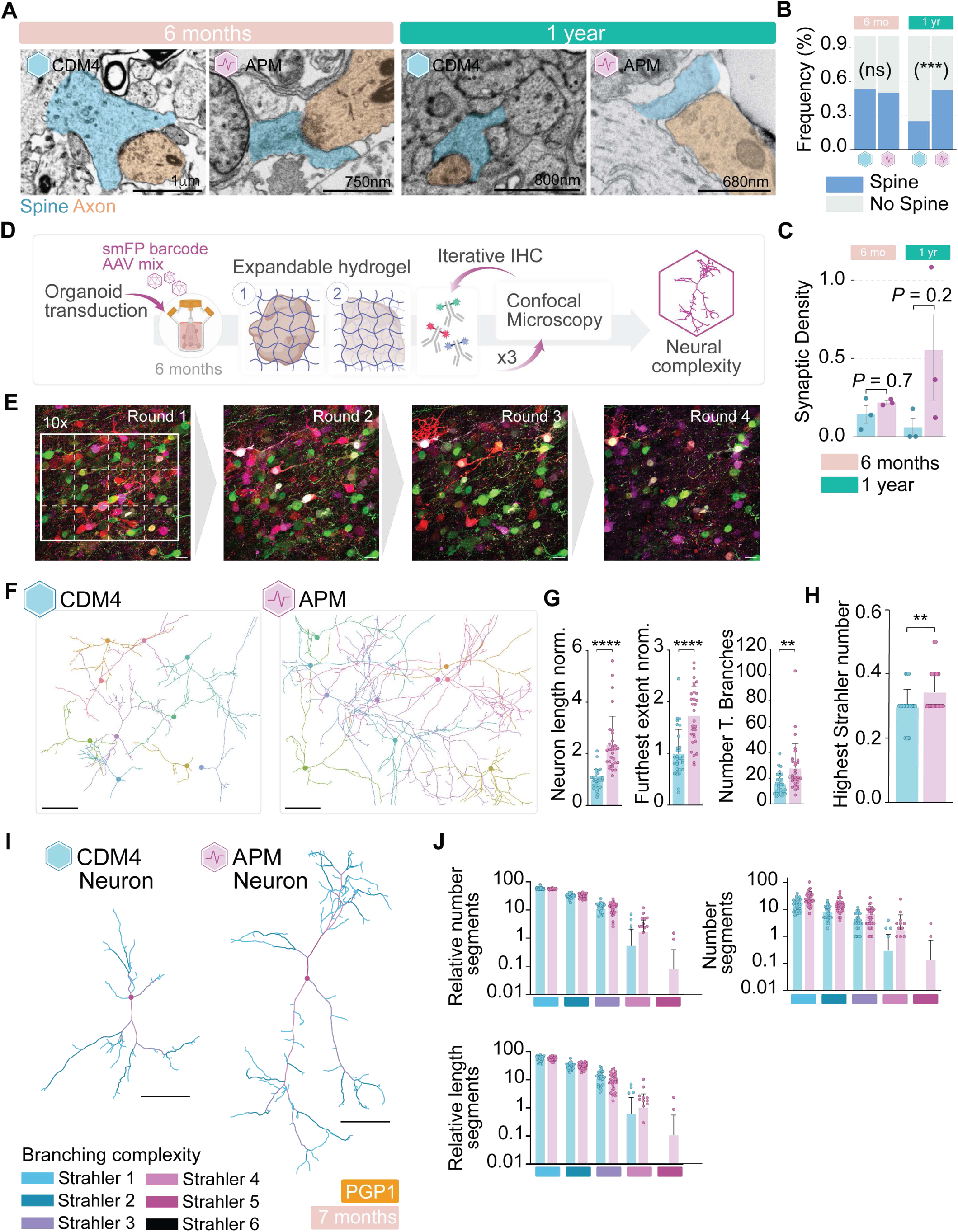
**A**, Representative electron microscopy images of the synaptic compartments. Scale bar: 1μm. **B**, Quantification of EM-Dendritic spine frequencies. Stacked barplot showing the frequencies of having or not having dendritic spines under each treatment at 6 and 12 months. For each age, a two-sided Fisher’s exact test was applied on the raw count data to indicate if there is a significant difference between CDM4 and APM in terms of having or not having dendritic spines (*P* values = 0.51 (6 months); 1.6 × 10^-4^ (12 months)). **C**, Quantification of EM-Synaptic density at 6 and 12 months. For each age, the measurement of synaptic density between CDM4 (n = 3) and APM (n = 3) was compared by the Wilcoxon rank-sum test (*P* values = 0.7 (6 months); 0.2 (12 months)). Bars represent the mean values with error bars indicating the standard error of the mean. **D,** Schematic of epitope tag barcoding and Expansion Microscopy workflow. **E**, Low-magnification overview of four rounds of iterative immunohistochemistry. Scale bar: 100 μm. White box indicates area for high-magnification tile scan imaging for neuronal tracing. **F**, Tracing and morphological reconstruction of ten representative individual neurons in CDM4 and APM conditions. Scale bar: 162.5 μm. **G**, Quantifications of different parameters related to length and complexity of neuronal arborization. Left: neuron length normalized to CDM4, unpaired t-test, *P* < 0.0001. Middle: furthest extent normalized to CDM4, unpaired t-test, *P* < 0.0001. Right: number of terminal branches, unpaired t-test, *P*= 0.0018. **H**, Comparison of highest Strahler number in CDM4 and APM conditions, unpaired t-test, *P* = 0.0034. **I,** Visualization of Strahler analysis for one representative CDM4 and APM neuron. Terminal branches receive Strahler number one. When two branches with the same Strahler number meet at a branch point, the Strahler number of the next segment increases by one. **J**, Quantification of absolute number, relative number, and relative length of different Strahler number segments in CDM4 and APM conditions.

To quantify synapse density across these large sections, we refined an existing deep learning algorithm(Pavarino et al., 2023) for the automatic detection of synapses in each section. While no statistically significant differences in synaptic density were observed between groups at either time point (Wilcoxon rank-sum test *P* values = 0.7 at 6 months, 0.2 at 12 months; **Fig. 3A, C**), at 12 months, synaptic density in 2 of 3 CDM4-treated organoids was nearly undetectable (0.0008 synapses/µm³), while the third had 0.17 synapses/µm³. In contrast, APM-treated organoids all exhibited higher synaptic densities, ranging from 0.17 to 1.08 synapses/µm³.

To further explore ultrastructural features associated with maturation, we classified a small subset of synapses (103 predicted synapses in each section) based on their postsynaptic target. Synapses were categorized as being located on dendritic spines (henceforth spines) if they were either clear, well-defined spines attached to a dendritic shaft, or putative spines—structures with morphology consistent with spines observed in 2D EM sections (**Fig. 3A, B**). Synapses that did not meet these criteria were classified as being located on non-spine elements. Our quantification revealed a significant increase of spine proportion in the APM-treated organoids at 12 months (**Fig. 3C**, 25% vs 52% in CDM4 and APM, respectively; two-sided Fisher’s exact test, *P* values = 0.51 (6 months); 1.6×10^-4^ (12 months)). Altogether, these findings suggest that APM treatment supports the maintenance of mature ultrastructural features in neurons over time. Notably, the higher proportion of synapses formed on dendritic spines in APM-treated organoids is consistent with enhanced synaptic maturation and stabilization, features that are critical for the long-term integrity of synaptic circuits.

We next asked whether neuronal complexity (i.e., arborization and process outgrowth), which is a clear measure of cortical neuron maturation *in vivo*, was enhanced by culture in APM. We adapted epitope tag barcoding(Dhainaut et al., 2022; Kudo et al., 2022; Rovira-Clavé et al., 2022; Wroblewska et al., 2018) as a novel tracing strategy(Park et al., 2025) to reconstruct the morphology of individual neurons in densely packed organoid tissue (**Fig. 3D, E and Supplementary Fig. 4E**). This method leverages stochastic expression of distinct epitope tag combinations in the form of antigenically orthogonal spaghetti monster fluorescent proteins (smFPs)(Viswanathan et al., 2015) to ensure robust cytosolic barcode filling (see **Methods**). Combined with the Magnify Expansion Microscopy strategy(Klimas et al., 2023a), optimized for robust multi-round epitope tag read-out (see **Methods**), this approach enables high-resolution detection of individual neurons and tracing of their fine processes in densely labeled tissue.

We infected organoids with an adeno-associated virus (AAV) mix for three weeks (n=3 organoids per condition, at 7 months in culture). Each of the nine AAVs in the mix contained a plasmid for the expression of an smFP, with eight copies of either V5, Myc, Flag, Ollas, E, S, HSV, Protein C or Ty1 epitope tag, and performed iterative post-expansion immunostaining of the expanded organoid sections to read out epitope tag expression and reconstruct 30 individual neurons for each condition (**Fig. 3D, E and Supplementary Fig. 4E, F**). APM-treated neurons exhibited significantly greater total neurite length, longer maximal process extension, and a higher number of terminal branches, revealing that both increased branching behavior and greater process outgrowth into the tissue contribute to enhanced neuronal complexity under APM culture (**Fig. 3F, G**). Strahler order analysis further showed that APM neurons achieved significantly higher maximum Strahler numbers, confirming greater complexity of neuronal arborization (**Fig. 3H, I)**. Notably, the distribution of lower Strahler indices remained comparable between conditions, indicating that increased arborization in APM organoids still follows the same principles of hierarchical neuronal architecture (**Fig. 3J)**.

Together, these results demonstrate that APM culture conditions sustain organoid activity over extended culture and promote synapse development and the acquisition of neuronal morphological complexity.

To determine whether morphological and functional maturation were reflected in changes at the transcriptional level (**Fig. 4, Supplementary Fig. 5 and 6**), we performed single-cell RNA sequencing across organoids derived from 4 different donors and collected from 6 to 12 months in culture (6 months: CDM4: n = 10, 64,817 cells; APM: n = 9, 41,231 cells; 9 months: CDM4: n = 6, 18,104 cells; APM: n = 3, 8533 cells; 12 months: CDM4: n = 4, 7802 cells; APM: n = 3, 6084 cells).

**Figure 4.**
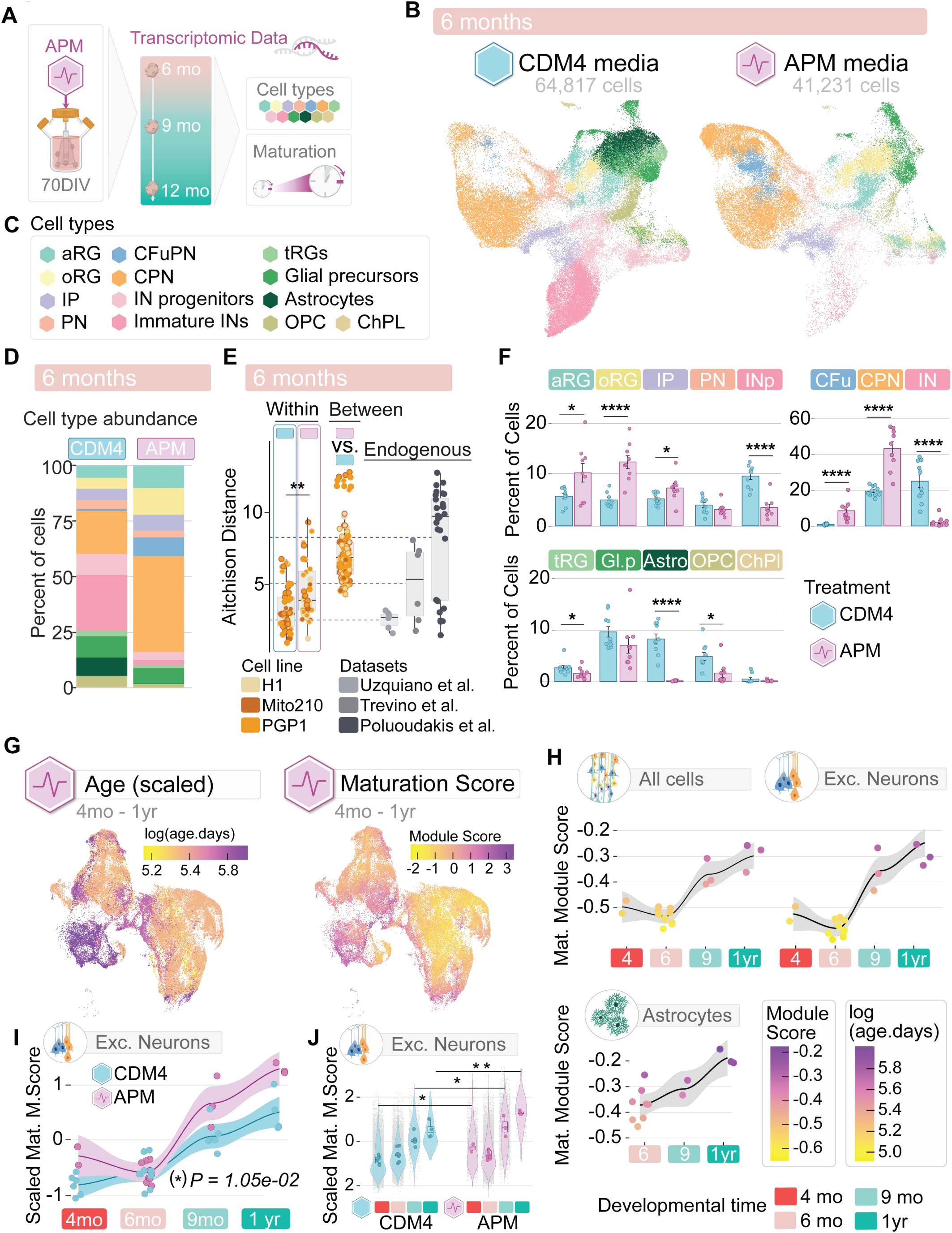
**A**, Schematics. **B**, UMAP of integrated scRNA-seq data from 6-month CDM4- (n = 10 organoids, 64,817 cells) and APM-treated (n = 9 organoids, 41,231 cells) organoids, colored by cell types. **C**, The collection of annotated cell types. **D**, Stacked barplot showing the summarized cell type proportions under each treatment. **E**, Aitchison distance measuring the differences in cell type compositions at 6 months was calculated for unique pairs of samples within each treatment and between 2 treatment conditions. Aitchison distances between replicates within each treatment were compared by the Wilcoxon rank-sum test (*P* = 1.21 × 10^-3^). The means for Aitchison distances (dotted lines) for three different datasets of endogenous human fetal cortex were added. **F**, Changes of cell type proportions between CDM4- (n = 10) and APM-treated (n = 9) organoids at 6 months. Bars represent the mean percentage of cell types for each treatment with error bars indicating the standard error of the mean. The significance of cell type proportion differences was calculated by the Likelihood Ratio Test comparing negative binomial models with and without treatment as a fixed effect using anova function. Each model takes the log of the total number of cells per organoid as an offset, the genetic background of the organoid as a random effect. FDR-adjusted *P*-values are indicated by asterisks: *P* < 0.05 (*); *P* < 0.01 (**); *P* < 0.001 (***); *P* < 0.0001 (****). **G**, UMAP of merged scRNA-seq data of APM-treated organoids from 4 to 12 months (n = 17 organoids; 60,427 cells). Left: colored by log10(age in days); Right: colored by Z-scored DIALOGUE Maturation Score. **H**, Age-dependent trends in DIALOGUE Maturation Score along developmental time points in APM-treated organoids. Scores were averaged across cells either globally or by cell populations (Astrocytes: Astrocytes; Exc. Neurons: CFuPN, CPN, and PN) for each organoid, represented by jittered dots. R’s geom_smooth function was used to add smoothed conditional means (black lines) with default 95% confidence intervals (grey regions). **I**, In excitatory neurons, APM-treated organoids show increased DIALOGUE Maturation Scores in a time-series manner (CDM4: n = 29,452 cells from 26 organoids; APM: 27,337 cells from 17 organoids). The Maturation Scores were Z-scored on the entire dataset and then averaged across excitatory neurons per organoid, denoted by dots with colors representing each treatment. A baseline linear model was created to evaluate how scores change with age (score ∼ age). A second linear model was created incorporating treatment in addition (score ∼ treatment + age). The additive effect of Treatment was found to significantly improve model fit, as indicated by a one-sided F-test (*P* = 1.05 × 10^-2^). **J**, In excitatory neurons, APM-treated organoids show significantly higher DIALOGUE Maturation Scores at 4, 9, and 12 months. The grey dots in the background denote scores of individual excitatory neurons present in the data and violin plots show the distribution. The colored dots are added to denote averaged scores across cells per organoid to enhance visibility. Boxes are overlaid to summarize the distribution of scores on the organoid level. Linear mixed-effects models were used to model the scores by including which organoid the cell belongs to and the genetic background of that organoid as random effects (score ∼ (1|Organoid) + (1|Genotype)) and a weight factor to adjust for cell count differences in excitatory neurons across organoids. Models with or without treatment as an additive fixed effect were evaluated using the anova function, with significance determined by *P*-values derived from a likelihood ratio test (4 months: 2.05 × 10^-2^; 6 months: 5.48 × 10^-1^; 9 months: 1.65 × 10^-2^; 12 months: 2.03 × 10^-3^).

Reproducibility, assessed using Atchison distances(Antón-Bolaños et al., 2024), demonstrated that both CDM4 and APM-treated organoids clustered closely; at all ages in both conditions, inter-sample variability fell within the limits of variation found in endogenous tissue, indicating that APM cultures preserve the high reproducibility we previously demonstrated in this organoid model(Antón-Bolaños et al., 2024; Uzquiano et al., 2022) (**Fig. 4E, Supplementary Fig. 6G**).

Following our hypothesis that the greater neuronal activity promoted by APM would favor maintenance of excitatory neuronal populations over extended culture times, we examined the proportions of individual cell types in each condition over time. We were particularly interested in testing for prolonged maintenance of callosal projection neurons (CPNs), a late-born population of excitatory neurons that are expanded in humans compared to other mammals and are associated with the evolution of higher cognitive functions(Berg et al., 2021).

Notably, as early as 6 months (CDM4: n = 10, 64,817 cells; APM: n = 9, 41,231 cells), we observed a marked difference in cellular composition between media conditions: the proportions of excitatory neuron subtypes—including callosal projection neurons (CPNs) and corticofugal projection neurons (CFuPNs)—were increased in APM, while glial populations and inhibitory neurons were reduced (FDR-adjusted *P* values: 1.15 × 10^-2^ (aRG), 3.38 × 10^-7^ (oRG), 1.95 × 10^-2^ (IP), 9.23 ×10^-8^ (CFuPN), 2.69 × 10^-8^ (CPN), 1.24 × 10^-5^ (IN progenitors), 4.19 × 10^-6^ (Immature IN), 4.81 × 10^-2^ (tRG), 1.52 × 10^-1^ (Glial precursors), 4.33 × 10^-13^ (Astrocytes), 2.46 × 10^-2^ (OPC), negative binomial mixed-effect (NBME) model, **Fig. 4B-D, F**). Notably, the enrichment of CPNs was maintained at 9 and 12 months [FDR-adjusted p values: 7.05 × 10^-3^ (9-month CPNs) and 1.96 × 10^-2^ (12-month CPNs)]. Together, this indicates that APM culture conditions selectively favor the long-term maintenance and stability of excitatory cortical neurons, including CPNs (**Supplementary Fig. 6F**).

To evaluate whether APM treatment promoted transcriptional maturation as well as maintenance of neuronal populations, we applied the DIALOGUE-based module scoring approach described in **Fig. 1D**, using the same maturation-associated gene modules derived from the Velmeshev et al. dataset of endogenous brain(Velmeshev et al., 2023). As in the CDM4 organoids, APM organoids showed strong correlation between time in culture and maturation module score (**Fig. 4G, H**). Importantly, this approach enabled a direct comparison between APM- and CDM4-cultured organoids along the temporal trajectory. For excitatory neurons (**Fig. 4J**), across time, APM organoids scored consistently higher in maturation index compared to CDM4 organoids (one-sided F test *P* = 1.05× 10^-2^, linear models); these differences were significant at all time points except at 6 months (likelihood ratio test *P* values = 4-month: 2.05 × 10^-2^; 6-month: 5.48×10^-2^; 9-month: 1.65×10^-2^; 12-month: 2.03×10^-3^; linear mixed-effects models) (**Fig. 4J**).

Taken together, these results show that promoting neuronal activity through culture in activity-permissive media results in neurons that display greater morphological and transcriptional maturity compared to those from organoids cultured in conventional media, while still being generated according to endogenous molecular programs of development. Notably, our ability to maintain and profile excitatory neurons over these extended culture periods has enabled the generation of a comprehensive multi-modal dataset, comprising molecular, morphological, and electrophysiological data on human excitatory cortical neurons at an unprecedented, advanced stage of *in vitro* development.

### Cortical progenitors record and retain memory of developmental time

Elegant work in the developing mouse and chick embryos has shown that developing cells track developmental time, progressively restricting their fate potential to execute temporally-appropriate developmental programs even if transplanted into a heterochronic host(Oberst et al., 2019; Shen et al., 2006). To test whether cells in older organoids retain a “memory” of the time they had spent in culture, we applied a functional measurement of aging in the dish. We probed cells derived from organoids of different ages for distinctions in fate potential, manifested as their ability to generate early versus late progeny. To this end, we leveraged protocols to produce chimeric organoids (“chimeroids”(Antón-Bolaños et al., 2024)) from either a mix of old cells (collected at 9-12 months), or a mix of old (9-12 months) and young (15DIV) cells.

First, we produced mono-chronic chimeroids (MonChr) by dissociating and re-mixing cells from late-stage cortical organoids (9 to 12 months old). Despite their prolonged time in culture, these cells could be dissociated and reaggregated to initiate the formation of a new organoid, although the resulting organoids were smaller, likely due to the limited proliferative potential of the old neural progenitors (**Fig. 5A, B**). To examine the cell composition of the resulting MonChr, we profiled pooled chimeroids (combining 15–20 chimeroids per batch) using scRNA-seq at 15–19 DIV post-reaggregation.

**Figure 5.**
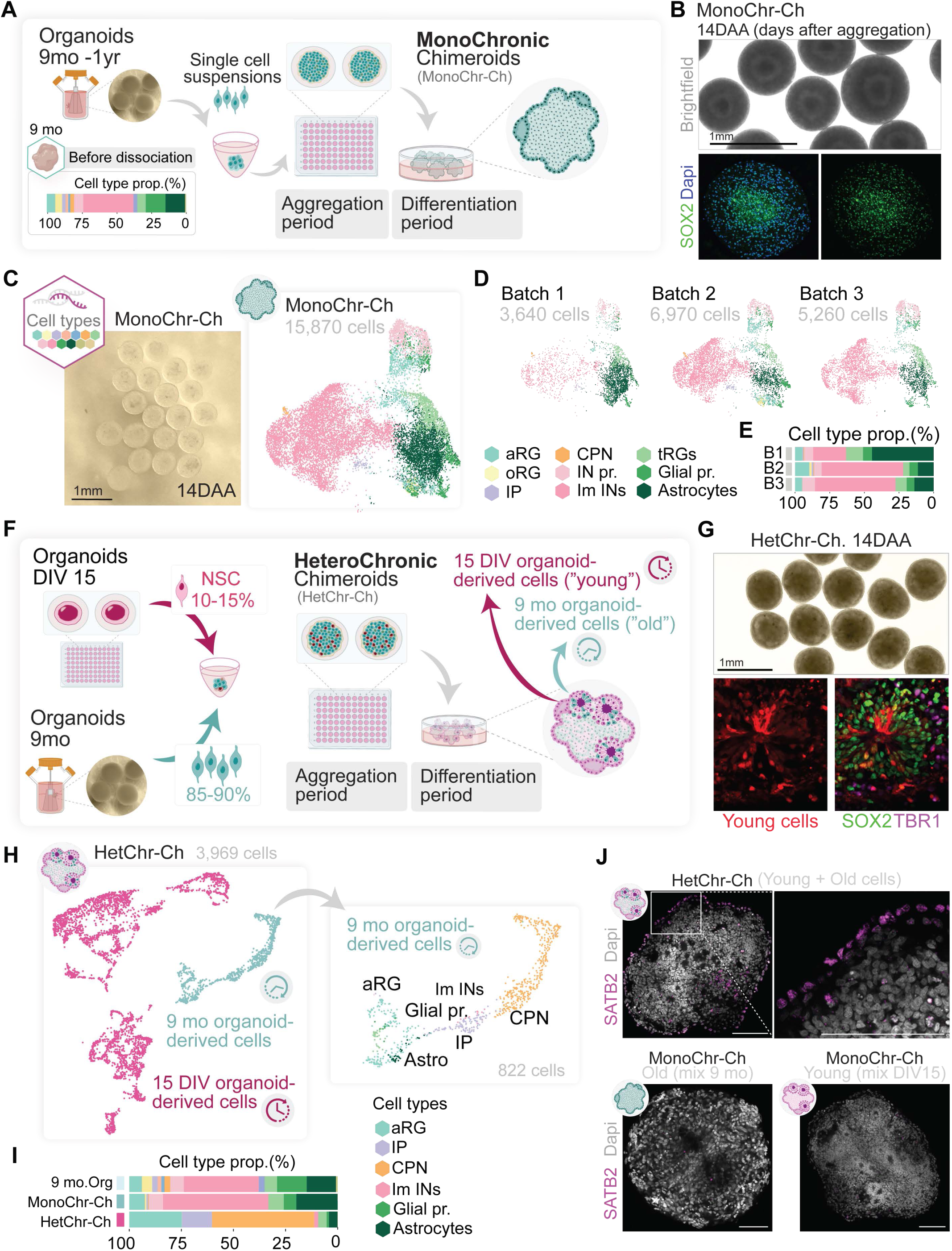
**A**, Schematic of MonoChronic culture generation, including a stacked bar plot (bottom) showing the cell type composition of 9-month organoids. **B**, Representative brightfield image of Monochronic Chimeroids (MonChr-Ch) and relative immunostaining. **C**, UMAP plot showing cells from MonChr-Ch (n=15,870 cells from 3 Chimeroids), colored by cell type. **D,** UMAP of MonChr-Ch split by each sample. **E**, Stacked barplot showing the summarized cell type proportions for each MonChr-Ch sample **F**, Schematic of HeteroChronic culture generation. **G**, Representative brightfield image of Heterochronic Chimeroids (HetChr-Ch) and relative immunostaining. **H**, UMAP plot showing cells from a HetChr-Ch sample (n=3,969 cells), colored by temporal groups of cells and cell type annotation of the old counterpart. **I**, Stacked barplot shows the relative proportion of cell types within the canonical 9-month organoids, the 9-month MonChr-Ch, and the 9-month-old portion of the HetChr-Ch. **J**, SATB2 immunostaining across MonChr-Ch and HetChr-Ch conditions. Scale bar: 100 μm.

We found that the MonChr samples mirrored the cellular composition expected for organoids grown for 9 months of culture, with all major cell types (>5% abundance in 9mo organoids) present. Of these major cell types, the MonChr had reduced prevalence of glial precursors (negative binomial GLMEM, P = 8.5 × 10-4) and oRGs (P =0.038); abundances of immature INs, IN progenitors, tRGs, aRGs, and astrocytes were not significantly different between MonChr and 9mo organoids (P > 0.1), despite having developed for only 15 days post reaggregation (**Fig. 5A, C-D**). This suggests that old progenitors retain a memory of the time they had previously spent in culture, producing progeny that is appropriate for their age (9-12 months). It is possible that this outcome may be the result of a lack of signals instructive of earlier fates within the old MonChr, as well as the possibility that the progeny may have been generated before dissociation and remixing; however, we noted that the chimeroids increased in size between initial reaggregation and collection, indicating that some proportion of cells in the final chimeroids were newly-generated through proliferation.

To probe the system further, we tested the ability of old cells to respond to instructive conditions of early cell fates produced via co-culture with cells from younger organoids (Heterochronic chimeroids, or HetChr). For this, progenitors were collected from late-stage cortical organoids (9 to 12 months old) and mixed with developmentally asynchronous progenitors derived from young organoids, collected at 15 DIV. HetChr were allowed to develop and profiled by scRNA-seq at 15 DIV after aggregation (**Fig. 5F, G)**. We employed different donor cell lines for young (PGP1) and old (GM) organoids; this strategy allowed us to demultiplex donor identity at the single-cell level and resolve the donors (and thus age) contributing to each differentiated cell type in the mixed organoid (see **Methods**(Antón-Bolaños et al., 2024; Mitchell et al., 2020)). To examine the effects of the heterochronic culture on the 9mo-derived cells (hereafter defined as ‘old-derived’), we subset the data from the HetChr samples to examine only all cells derived from the ‘old’ organoids. In contrast to their fate in monochronic chimeroids, derivatives of these ‘old’ cells now exhibited a notable shift in cell-type composition compared to standard organoids at 9-12 months (**Fig. 5H-I**). Specifically, old-derived cells in HetChr chimeroids showed a strikingly higher proportion of callosal projection neurons (CPNs; 49.0%), compared to both the MonChr group (Rep1 = 0.55%, Rep2 = 0.50%, Rep3 (11mo) = 0.02%), and the 9-month standard organoids (CPNs; 1.1%). We performed immunostaining to assess the expression of SATB2 protein (a marker of CPNs) in young MonChr organoids (1-month old, 15 DIV after reaggregation), old MonChr organoids (9-month old, 15 DIV after reaggregation), and HetCh organoids (9-month and 1-month old, 15 DIV after reaggregation), all cultured in CDM4 medium. In line with the scRNA-seq results, SATB2 signal—absent in both old and young MonChr cultures—was clearly detected in the HetChr condition (**Fig. 5J** and **data not shown**). In contrast, young-derived cells produce the expected, early, cell types at DIV 30 (15 DIV + 15 DIV after aggregation, **Supplementary Fig. 7A**). These results indicate that old progenitors are still plastic after 9 months in culture and can respond to instructive neurogenic signals to make neurons again; however, they have recorded the passage of time and retain a memory of the developmental steps already performed, as demonstrated by the fact that in only 2 weeks they can generate late stage neurons (CPNs), while their young progenitor counterparts undergo differentiation into much earlier cell fates within the same chimeric organoid.

To confirm these findings, we repeated the same experiment but using progenitors derived from the embryonic mouse brain. This system gives the distinct advantage of fast development and endogenous origin of the progenitor cells. We dissected the cerebral cortex from E11.5 mouse embryos and seeded single cells in suspension to generate endogenous mouse cortical chimeroids (**Supplementary Fig. 7B-D)**. After two weeks in culture, these organoids predominantly expressed astrocytic markers as well as the CPN marker SATB2 (**Supplementary Fig. 7E)**. We then created chimeroids by either mixing progenitors from the two-week-old organoids alone (MonChr), or mixing cells from the two-week-old organoids with freshly dissociated, young, E11.5 cells (HetChr, **Supplementary Fig. 7F**). We analyzed the cultures 1 day after aggregation, to accommodate the faster developmental timeline of the mouse compared to humans (**Supplementary Fig. 7G**). As observed in the human model, while the old MonChr mouse cultures showed minimal SATB2 expression, the HetChr condition exhibited a significantly higher proportion of SATB2 positive cells (ANOVA *P*-value < 0.05). Notably, control monochronic chimeroids containing only young cells took longer to begin to produce SATB2 positive cells, in line with the fact that they had not yet completed earlier developmental steps. The data indicated that the cells in the older organoids were specifically reactivating neurogenic programs suited to their later developmental age (**Supplementary Fig. 7G, H)**.

Together, these results indicate that cortical organoid cells retain a form of temporal memory. This is reflected in their distinct fate potential at late ages compared to young progenitors cultured for shorter times. Old progenitors continue to produce temporally-appropriate progeny after reaggregation and, even when exposed to signals instructive of an earlier neurogenic fate, they “skip” production of early progeny to generate neurons normally produced after months in culture.

## Discussion

The speed of development of cells and organs is markedly different across species(Casimir et al., 2024). The human brain in particular displays extremely long periods of neoteny, notoriously taking close to two decades to form and fully mature. Human brain organoids that can continue to develop over years, rather than weeks and months, offer a unique opportunity to experimentally access processes and mechanisms of human brain postnatal maturation that are species-specific and cannot be easily accessed, experimentally, *in vivo*.

Endogenous brain tissue typically exhibits epigenetic stability after early development, with DNA methylation patterns maintained across later postnatal stages(Luo et al., 2016). In contrast, most *in vitro* systems gradually show global methylation loss, erosion of regulatory regions, and increased epigenetic variability over time(Franzen et al., 2021). Our finding that organoids display global stability of DNA methylation patterns for five years in culture, point at a robust and resilient system that reproduces the epigenetic landscape of the endogenous brain. Notably, the observed methylation changes at loci associated with key cell lineage decisions, also points at a dynamic system that progressively attains developmental milestones. These findings demonstrate that temporally structured regulation of cortical development can unfold over unprecedented timescales, even outside the context of the body.

Activity is critically important for brain development and maturation. We show that in media supportive of spontaneous activity, organoid can support neuronal health and promote their maturation past any previously known limits. These findings motivate future efforts to parallel integration of early spontaneous activity with later evoked activity, obtained via the incorporation in organoids of sensory input, for example by integrating cells of the retina and/or the cochlea.

As the brain develops, matures and ages, cells accumulate epigenetic changes that alter their function and potential. The fact that cells derived from old organoids, even when exposed to early instructive signals, largely generate late neuronal progeny, provides an important functional validation for the idea that cells in organoids do not just survive for years in culture, they progressively age, recording the passage of time in epigenetic changes that ultimately alter their function and fate potential. This opens the door to experimental investigation of the mechanisms of these largely inscrutable processes of brain maturation. In addition, the ability of cells derived from old organoids to skip steps of development that they have already performed, and to generate in two weeks late neuronal progeny that would normally take over two months to produce, is remarkable. In practical terms, this points at new strategies for rapid generation and more extensive maturation of neuronal populations such as Callosal Projection Neurons (CPNs), which are some of the most evolutionarily divergent neurons of the human neocortex, associated with the acquisition of uniquely human cognitive functions, yet are born late and thus are difficult to form and mature in organoids.

Altogether, organoids cultured over extended timelines and the multi-modal wealth of data produced from them represent both a powerful experimental system and the information to fuel understanding of the largely unexplored mechanisms governing human brain neoteny, maturation, and evolution.

## Acknowledgements

We thank Juliana R. Brown (from the P.A. laboratory) for input and assistance in editing the manuscript, Nalini Oliver, Aiman Altaf, Lila L. Lyons, Noah Kozub and Sarah Tropp for excellent technical assistance and all the members of the Arlotta laboratory for discussions; B. Cohen (McLean Hospital) for the Mito210 iPSC line; G. Church (Harvard University) for the PGP1 iPSC line; M. Talkowski (MGH) for the GM08330 iPSC line; the Broad Genomics Platform for sequencing. We thank Mansour Alawi for helping D.L with the expansion microscopy analyses. We thank Sam Rodriques and Andrew Payne for their contribution to early development of the epitope tag barcoding constructs. This work was supported by grants from the Stanley Center for Psychiatric Research to P.A., and J.Z.L., the Broad Institute of MIT and Harvard, the National Institutes of Health (P50-MH094271 and RF1-MH123977 to P.A., R01-MH112940 to P.A. and J.Z.L), and the Klarman Cell Observatory (A.R.).

## Authors’ contribution

N.A.B, I.F and P.A. conceived and designed the experiments. N.A.B, I.F and, S.A generated, cultured, and characterized all the organoids in this study, with the help of S.S, and P.A. supervised their work and contributed to data interpretation.

N.A.B and I.F worked on cell type assignments and scRNA-seq data analysis, and A.W. and T.F performed all the computational work under supervision by A.R and P.A

A.S.K., performed methylome experiments and helped with analysis and interpretation. A.K. prepared the methylome libraries for sequencing. M.S performed the computational analyses for the methylome data under the supervision of H. K. and A.M.

N.A.B., I.F., S.A and S.S. performed IHC and image acquisition. R.K. performed MEA recordings and Y.S.M performed data analysis with the help of R.K. M.M.C performed all the electron microscopy experiments and ground truth analysis with the help of S.A and Y.M. Y.M worked on the quantification algorithm for synapses quantification under the supervision of J.L. N.A.B and I.F performed scRNA-seq experiments with help from X.A and supervision of P.A. and J.Z.L.

D.L. designed and performed the expansion microscopy epitope tag barcoding experiments and analyses, with help from E.Z, and B.A. contributed to early development of the epitope tag barcoding constructs, all under the supervision of E.S.B.

N.A.B and A.S.K performed the mouse experiments. N.A.B. and I.F performed scRNA-seq library preparation for the vast majority of the preparations included in this study.

N.A.B, I.F. and P.A. wrote the manuscript with contributions from all authors. All authors read and approved the final manuscript.

## Materials & Correspondence

Correspondence and request for materials should be addressed to P.A.

## Code Availability

Code used during data analysis is available at [available before publication].

## Data Availability

Read-level data from scRNA-seq have been deposited in a controlled access repository at [available before publication] while count-level data and meta-data have been deposited at the [available before publication]. Any additional information required to reanalyze the data reported in this paper is available from the lead contact upon request. Data from previous publications that were used in this study can be found at [available before publication].

## METHODS

### Ethics statement

All experiments involving human cell lines were approved by the Harvard University IRB and ESCRO committees. All animal experiments were conducted according to protocols approved by the Institutional Animal Care and Use Committee (IACUC) of Harvard University. All experiments were performed in accordance with relevant guidelines and regulations, and in accordance with informed consent obtained from the donors of the originating cells or tissues.

### Human pluripotent stem cell culture

All human pluripotent stem cell (PSC) lines were maintained as previously detailed(Antón-Bolaños et al., 2024; Velasco et al., 2019). Briefly, MTESR1 medium (StemCell Technologies), mTESR^+^ medium (StemCell Technologies), or StemFlex medium (Gibco), all with 1% of added Penicillin-Streptomycin Solution (Corning), were employed for culture of stem cells in cell culture dishes (Falcon) pre-coated with 1% Geltrex (Gibco), at 37°C in 5% CO2. All human PSCs were maintained below passage 55, tested negative for mycoplasma (assayed with MycoAlert PLUS Mycoplasma Detection Kit, Lonza) and were karyotypically normal (G-banded karyotype test performed by WiCell Research Institute).

### Characterization of the PSC lines

The psychiatric control Mito210 male induced PSC line was provided by B. Cohen (McLean Hospital); the PGP1 male induced PSC line was provided by G. Church (Harvard University); the GM08330 male induced PSC line was provided by M. Talkowski (MGH) and was originally derived from fibroblasts obtained from the Coriell Institute for Medical Research(Paulsen et al., 2022; Uzquiano et al., 2022; Velasco et al., 2019). The H1 male human embryonic stem cell line (also known as WA01) was purchased from WiCell; and the 11a induced PSC line was obtained from the Harvard Stem Cell Institute. The female CW50037 induced PSC line was from the California Institute for Regenerative Medicine (CIRM) iPSC collection. All lines were authenticated as follows: The PGP1 induced PSC line was authenticated by short tandem repeat (STR) analysis (performed by TRIPath). The Mito210 induced PSC line was authenticated with genotyping analysis (Fluidigm FPV5 chip) performed by the Broad Institute Genomics Platform. The CW50037 iPSC line was authenticated using SNP genotyping (Illumina Global Screening Array (GSA) from Illumina; processed at the Genomics Platform, Broad Institute) for cell line identification and detection of chromosomal abnormalities. The H1 and GM08330 lines were authenticated by STR analysis (performed by WiCell). The GM08330 parental line has a previously-reported(Paulsen et al., 2022) interstitial duplication in the long (q) arm of chromosome 20; all other lines were karyotypically normal.

### Cortical organoid and chimeroid differentiation

Dorsally patterned cortical organoids were generated following a protocol previously published by our group(Velasco et al., 2019). Briefly, on day 0, feeder-free cultured human PSCs, 75–85% confluent, were enzymatically dissociated to single cells with Accutase (Gibco), and 9,000 cells per well were reaggregated in ultra-low cell-adhesion 96-well plates with V-bottomed conical wells (sBio PrimeSurface plate; Sumitomo Bakelite) using the same pluripotent cells medium in which they were previously maintained. At day 1, 80 μl of media was replaced with Cortical Differentiation Medium (CDM) I, containing Glasgow-MEM (Gibco), 20% Knockout Serum Replacement (Gibco), 0.1 mM Minimum Essential Medium non-essential amino acids (MEM-NEAA) (Gibco), 1 mM pyruvate (Gibco), 0.1 mM 2-mercaptoethanol (Gibco), with 1% of Penicillin-Streptomycin Solution (Corning). From day 0 to day 6, ROCK inhibitor Y-27632 (Millipore) was added to the medium at a final concentration of 20 μM. Patterning small molecules, Wnt inhibitor IWR1 (Calbiochem) and TGFβ inhibitor SB431542 (Stem Cell Technologies), were added from day 0 to day 18, at a concentration of 3 μM and 5 μM, respectively.

Single donor and multidonor Chimeroids were generated as previously described(Antón-Bolaños et al., 2024). Briefly, patterned EBs between day 15-18 were dissociated into single-cell suspensions using a modified papain-based protocol(Antón-Bolaños et al., 2024). After confirming morphology, organoids were enzymatically and mechanically dissociated, then filtered and resuspended in CDM1 medium with ROCK inhibitor. Cell suspensions from different donors (in the case of multidonor chimeroids) were mixed in equal ratios and reaggregated (18,000–20,000 cells/well) in ultra-low-adhesion 96-well plates. Two days later, embryoid bodies were transferred to low-attachment dishes and cultured under orbital agitation. Medium was sequentially changed to CDM2, CDM3 (at DIV35), and CDM4 (at DIV70) as per established protocols(Velasco et al., 2019). For APM, organoids at day 70 were transferred into a medium containing BrainPhys Basal (Stem Cell Technologies), 20ng/mL N2 Supplement (1x Thermo Fisher Scientific), 20ng/mL B27 Supplement (1x) (Thermo Fisher Scientific), 1% Penicillin Streptomycin Solution 100X (Corning), 1% Minimum Essential Medium non-essential amino acids (MEM-NEAA) (Gibco), 1% Glutamax (Gibco), and Amphotericin B (Thermo Fischer Scientific). After filtration, GDNF and BDNF at a final concentration of 20 ng/mL were added. Along with dCAMP (Milliporesigma), Ascorbic Acid (StemCell Technologies), and Laminin (Thermo Fischer Scientific) at a concentration of 1mM, 200nM, and 1ug/mL. Half of the medium was replaced with fresh medium twice per week. HeteroChronic and MonoChronic cultures were obtained using the previously described chimeroid strategy; however, papain dissociation time and mechanical dissociation were adjusted based on organoid age (45 minutes for older organoids that were previously minced with a blade, and 25 minutes for younger organoids). While the chimeroid strategy is extremely efficient in younger organoids, the success rate of the heterochronic cultures remains low (approximately 1 in 6). This is likely due to the dual requirement for sufficient patterning in the young organoids and adequate stemness in the older cells, both of which are necessary to ensure robust signaling from the young tissue and sufficient recovery of older cells for analysis at day 14 post-mixing.

### Mouse experiments

We used wild-type CD1 and C57Bl/6 mice (Charles River Laboratories). Animals were scarified to obtain embryos and staged carefully according to Theiler stages for E11.5 and the neocortex was dissected. The same dissociation protocol described for the human chimeroids has been used with some modifications. Dissociated progenitors were seeded to aggregate over night at the density of 12,000 cells per well. The next day, the media was switched to CDM2 for three days. On day 4 after seeding, the media were changed to CDM3 and CDM4 on day 6. We quantified the number of nuclei with positive SATB2 expression across three conditional groups: Mon-Ch (old), Mon-Ch (young), and Het-Ch from cultures 1 day (D1AM; for each group: n = 3 replicate organoids × 2 batches) and 2.5 days (D2.5AM; for each group: n = 3 replicate organoids × 1 batch) after mixing, respectively. For evaluating the overall effect of condition on the proportion of nuclei with positive SABT2 expression at D1AM, binomial generalized linear models (GLM) were used to model the number of SATB2-positive nuclei out of total nuclei per organoid and adjust for batch by including batch as a fixed effect. Models with or without adding a condition as a fixed effect were evaluated by a likelihood ratio test using the anova function. Following the observation of the global effect of the condition being statistically significant (ANOVA *P*-value < 0.05), estimated marginal means for each group were derived from the fitted full binomial GLM by using the R package emmeans, and Tukey-adjusted pairwise comparisons were performed between groups. For D2.5AM, due to the absence of more than one batch level to adjust for, we proceed with computing estimated marginal means and performing pairwise comparisons between groups.

### Fixation and processing of samples for cryosectioning

Organoids were fixed in 4% paraformaldehyde (PFA) (Electron Microscopy Services) overnight (O/N) in a 12-well plate (Falcon) at 4°C, washed with 1X phosphate buffered saline (PBS) (Gibco) 3 times, and cryoprotected in a 30% sucrose solution (Sigma) in PBS (O/N) at 4°C. Gelatin solution containing 10% of bovine gelatin (Sigma) and 7.5 % of sucrose (Sigma) was prewarmed at 37°C for 15 min. The 30% sucrose solution was removed from the samples and exchanged for the prewarmed gelatin, and the samples were incubated at 37°C for 15 min. Meanwhile, plastic molds were coated with a 2mm layer of warm gelatin solution and left to polymerize at room temperature (RT). The samples were then transferred to the pretreated plastic molds, and 1 mL of warm gelatin solution was added on top. Following polymerization at room temperature (RT) for 3 min, samples were pre-chilled at 4°C for 15-20 min. Finally, the molds were frozen in a cold bath containing 100% EtOH and dry ice for 2-3 min, and stored at -80°C indefinitely.

### Immunohistochemistry

For immunohistochemistry, 14 to 18 μm thick sections were cut using a cryostat (Leica). Cryosections were stabilized at RT for 5 min and blocked with 10% donkey serum (Sigma) + 0.3% Triton X-100 (Sigma) in PBS. Primary antibodies were diluted in the blocking solution and incubated O/N. After 4 washes with PBS, cryosections were incubated at RT with secondary antibodies diluted in PBS (1:1000) for 1 hour at RT, washed 4x with PBS, and stained with DAPI (1:10,000 in PBS + 0.1% Tween-20) for 5 minutes to visualize cell nuclei.

### Whole organoid immunofluorescence

Organoids were washed in wash buffer (PBS with 0.4% BSA) and fixed at RT for 30 minutes with 4% PFA. After fixation, organoids were stored in 1X PBS with 0.1% Tween-20 (P9416, Sigma). Permeabilization, blocking, and staining were performed as previously described(Grosswendt et al., 2020). Nuclei were counterstained with 0.5μg/mL DAPI for 30 minutes. Organoids were optically cleared using RIMS and mounted as previously described(Grosswendt et al., 2020; Sampath Kumar et al., 2023). Images were acquired with LSM880 (Zeiss) and LSM900 (Zeiss) using a 40X objective. Images were processed with ImageJ (version 2.14.0/1.54f). Nuclei were segmented and quantified using ZEN blue 3.1 Image Analysis and Intellesis software packages. Image segmentation was achieved using a previously trained ZEN Intellesis algorithm(Frey et al., 2012). SATB2 intensity was thresholded to define positive nuclei and counted across the various z-stacks of individual organoids.

### Microscopy

Immunofluorescence images were acquired using the Zeiss Axio Imager.Z2. Images on the Axio Imager.Z2 were acquired with a ×20 objective (pixel size, 0.325 μm) using the Apotome optical sectioning function. The Zen Blue software was used to perform tile stitching and apotome deconvolution before exporting the images. For the LSM900, z stacks with 3 μm step size were acquired with a ×20 objective (pixel size, 0.62 μm), followed by tile stitching using the Zen Blue software. Further processing, such as z projections, channel merging and adding scale bars, was performed in Fiji43. Images were adjusted for brightness and contrast; all adjustments were applied to the whole image.

### DNA extraction and methylome analyses

#### Genomic DNA extraction and whole-genome bisulfite sequencing

Genomic DNA was isolated from organoids using a previously described method(Haggerty et al., 2021). Briefly, organoids were thawed on ice and suspended in 500 μL of resuspension buffer (240 μL of Genomic lysis buffer (D4075, Zymo) + 20 μL of Proteinase K (D4075, Zymo) + 240 μL of nuclease-free water). The sample was thoroughly mixed, vortexed, and incubated at 55°C for 4 to 7 hours at 300 rpm. An equal volume of Phenol: Chloroform: Isoamyl alcohol (15593031, Thermo Fisher Scientific) was added, mixed by inversion, and then centrifuged at 13,000 g for 5 minutes. The aqueous phase (∼400 μL) was transferred to a new tube for precipitation. To this aqueous phase, 8 μL of Glycogen-Blue (AM9515, Thermo Fisher Scientific), 20 μL of 5M NaCl (AM9760G, Thermo Fisher Scientific), and 1 mL of Ethanol (E7023, Sigma) were added, mixed by inversion, and allowed to precipitate overnight at -20°C. The tubes were spun at 13,000 g for 45 minutes at 4°C. The supernatant was discarded, and 1 mL of 70% ethanol was added to the pellet, which was then spun down at 13,000 × g for 45 minutes at 4°C. The supernatant was discarded, and the pellet was air-dried for 10 minutes before being resuspended in elution buffer (low-EDTA TE pH 8, 15575020 Thermo Fisher Scientific) and incubated at 55°C for 10 minutes.

Genomic DNA was fragmented in a tube (MicroTube AFA Fiber pre-slit snap-cap 6x16mm (520045, Covaris)) using a Covaris S-series S2 Sonicator with the following settings: duty cycle, 10%; intensity, 5; cycles per burst, 200; maximum temperature, 7°C; duration, 76-90 seconds. Fragmented DNA was purified and concentrated using the DNA Clean & Concentrator kit (D4013, Zymo) and eluted in 22 µL of low TE (pH 8). 1 µL of the eluate was used for quality control on the Agilent TapeStation (HSD5000), resulting in an average fragment size of 237-322 bp. Bisulfite conversion was carried out using the EZ DNA Methylation-Gold kit (D5005, Zymo), with the final converted DNA eluted in 16 µL of low TE (pH 8). Libraries were prepared using the xGen-Methyl-Seq DNA Library prep kit (10009824, IDT), in combination with xGen UDI Primer Plate 2 (8 nt, 10009816, IDT). The number of PCR cycles used was 7. Each library underwent two final rounds of purification using AMPure XP beads (A63881, Beckman Coulter). Libraries were assessed for concentration, fragment size distribution, and primer-dimer absence using the Agilent TapeStation (HSD5000). Sequencing was conducted on an Illumina NovaSeq 6000 platform, generating 150 base paired-end reads.

#### WGBS processing

Raw reads were subjected to adapter and quality trimming using cutadapt(Martin, 2011) (v4.6; parameters: --quality-cutoff 20 --overlap 5 --minimum-length 25; Illumina TruSeq adapter clipped from both reads), followed by trimming of 10 and 5 nucleotides from the 5’ and 3’ end of the first read and 15 and 5 nucleotides from the 5’ and 3’ end of the second read. Trimmed reads were aligned to the human reference genome (hg19) using BSMAP(Xi and Li, 2009) (v2.90; parameters: -v 0.1 -s 16 -q 20 -w 100 -S 1 -u -R). Sorted BAM files were generated and indexed using samtools(Li et al., 2009) with the ‘sort’ and ‘index’ commands (v1.21). Duplicates were removed using the ‘MarkDuplicates’ command from GATK (v4.5.0.0; -- VALIDATION_STRINGENCY=LENIENT --REMOVE_DUPLICATES=true -- COMPRESSION_LEVEL 4 --ASSUME_SORT_ORDER coordinate)(McKenna et al., 2010). Methylation rates were called using mcall from the MOABS(Sun et al., 2014) package (v1.3.9.6; default parameters; --reportCpX A/C/T for non-CpG methylation calling). All methylation analyses were restricted to autosomes and only CpGs covered by at least 10 and at most 150 reads were considered for downstream analyses. Assessment of global, genome-wide methylation was performed using average (arithmetic mean) methylation levels. Region-specific methylation was assessed by calculating average methylation levels across features (bins, PMDs, HMDs, DMVs, repeats, super-enhancers) using bigWigAverageOverBed from UCSC tools for each sample, where a feature was only considered if at least three CpGs were covered within a region.

#### Differential methylation analysis

Differentially methylated regions (DMRs) were called using metilene(Jühling et al., 2016) (v0.2-8) between organoids cultured for 3 months and 5 years. DMRs were defined by an absolute minimum difference in methylation of 0.1 with a maximum distance of 300nt between CpGs within a DMR and a minimum of 10 CpGs per DMR and Bonferroni correction for multiple testing (parameters: –c 1 –d 0.1). DMRs were filtered to retain only those with a minimum absolute Pearson correlation coefficient of 0.6 between time in culture (in months) and mean DMR methylation rate across all time points. DMRs were classified as ‘Hypo DMRs’ and ‘Hyper DMRs’ based on the direction of change. DMRs were associated with CGIs and DMVs based on a minimum overlap of 1bp. DMRs were annotated to the nearest genes using GREAT(McLean et al., 2010) (v4.0.4; default parameters).

#### Genomic Feature Annotation

CpG islands were obtained from the UCSC Genome Browser (https://genome-euro.ucsc.edu/cgi-bin/hgTables). CpG shores were defined as 2 kb regions flanking CGIs upstream and downstream; shelves were defined as 2 kb regions flanking the shores.

DNA methylation valleys (DMVs) were defined based on WGBS data from the HUES64 human embryonic stem cell line(Charlton et al., 2020). To identify DMVs, a previously described sliding window approach was adapted(Hetzel et al., 2022). Briefly, 5 kb windows with a 1 kb step size were generated across autosomes using bedtools(Quinlan and Hall, 2010) makewindows. Average CpG methylation was calculated for each window, excluding CpGs located within CGIs, and separately for CGIs. Windows and CGIs with an average methylation below 0.15 and containing at least 10 CpGs were merged, excluding regions composed solely of CGIs.

Annotations of highly methylated domains (HMDs) and partially methylated domains (PMDs) were obtained from https://zwdzwd.github.io/pmd(Zhou et al., 2018b).

Repeat annotations were obtained from the hg19 UCSC RepeatMasker track, excluding entries on alternative haplotypes, fix patches, and chrMT.

#### Epigenetic clocks

CpG methylation values at low-coverage sites (coverage < 5) were imputed using the boostme package (v0.1.0). Imputation was performed on individual samples represented as objects from the bsseq package (v1.40.0). Imputed values exceeding one were capped at one. Array-based probe IDs were mapped to genomic coordinates using the Illumina EPIC array manifest (EPIC.hg19.manifest). The pan-tissue human methylation clock(Horvath, 2013) was applied using the DNAmAge function from the methylclock(Pelegí-Sisó et al., 2021) package (v1.10.0). Cortical DNA methylation age(Shireby et al., 2020) was estimated using the CorticalClock function based on the implementation provided at https://github.com/gemmashireby/CorticalClock. DNA methylation age was assessed by median absolute error (MAE) between time-in-culture of the organoids and predicted methylation age in months, as well as Pearson correlation between the two across all samples.

To assess mitotic history, DNA methylation at solo-WCGW CpGs, which are particularly susceptible to methylation loss with cell division(Zhou et al., 2018a), was quantified. Methylation was evaluated across three feature sets: all solo-WCGWs, solo-WCGWs within common PMDs, and solo-WCGWs in common PMDs represented on the HM450 array. Annotations were obtained from https://zwdzwd.github.io/pmd, and average methylation levels across feature sets were calculated per sample from non-imputed WGBS data.

#### CpA methylation

Forebrain super-enhancers were obtained from a previously published dataset of fetal human brain tissue and cerebral organoids(Luo et al., 2016). Mean CpA methylation levels at these regions were compared to background levels calculated across genome-wide 30 kb bins (approximating the average super-enhancer size), excluding bins overlapping with super-enhancers.

#### Visualizations

Unless stated otherwise, all statistics and plots were generated using R version v4.4.1. Violin plots were generated using the vioplot (v0.5.1) or ggplot2 (v3.5.2) package and show the kernel density estimation with embedded boxplots indicating the median, interquartile range, and whiskers extending to 1.5× the interquartile range. Circos plots were created using the RCircos (v1.2.2) package from tracks with averaged profiles over 50kb bins created using UCSC tools bigwigAverageOverBed. The differential signal was obtained by subtracting the 5-year profile from the 3-month profile. Cytoband and ideogram data for hg19 were obtained from the RCircos package. DMR methylation heatmaps and average tracks were created using the EnrichedHeatmap(Gu et al., 2018) package (v1.34.0) by applying the normalizeToMatrix function first, with 2500 bp extensions up- and downstream and a target ratio of 0.4. Pairwise genome-wide correlations of methylation rates between samples of consecutive time points were plotted using the smoothScatter function. Bar plots, scatter plots, and line plots were created using ggplot2.

### Electrophysiological analyses

Electrical signals were recorded using the Accura 3D CMOS-HD-MEA system by 3Brain (BioCAM DupleX system in combination with Accura 3D chips). The 3D chip contains 4,096 penetrating µNeedle electrodes with 90µm height arranged in 64 × 64 grid in a square measuring 3.8 × 3.8mm with a sampling rate of 20kHz and 12-bit resolution. CDM4 organoids were changed to APM media 14-21days before the analysis and the acute recording was performed at 37°C in a mini-incubator box filled with Carbogen on the intact organoid. BrainWave v5 was used to record spontaneous activity for 15-20mins. To test if the electrical activity measured was synaptic NMDA and AMPA receptor activity was blocked at the end of recordings using D-AP5 (at 150μM) and DNQX (at 30μM), respectively. Additionally, action potentials were blocked by bath application of TTX (1μM). Spike detection was carried out by Kilosort4(Pachitariu et al., 2024), followed by analysis of the spike train using custom MATLAB scripts (version 24.1, R2024b, The MathWorks, Inc.). The detection of rapid spiking periods (burst) was performed by implementing the Maximum Interval(Kirillov, n.d.) algorithm (Max. Interval = 170 ms, Max. End Interval = 400ms, Min. Interval Between Bursts = 800 ms, Min. Duration of Burst = 40 ms, Min.Number of Spikes = 4). The analysis of network bursting was performed on the basis of the population-averaged spiking rate along all detected units. A peak in the population signal was considered to be a network burst if it met the following criteria: (1) the peak amplitude was greater than 4 times the s.d. of the noise value; (2) a set of bursting cells composed of at least 20% of total cells were active during that population spike; and (3) a cell was considered part of the set of bursting cells only if it participated in at least 50% of the network bursts. The peaks of the network bursts were used to measure the inter-burst interval (IBI), and the network burst rate was obtained from the average IBI.

### Electron microscopy analyses

A total of twelve organoids (three organoids per time point and condition: two H1 and one 11a genetic background at 6 months, and one H1, one 11a, and one PGP1 genetic background at 12 months for both CDM4- and APM-treated conditions) were immersion-fixed in 2% paraformaldehyde (EMS, 15710), 2.5% glutaraldehyde (EMS, 16220) and 0.003M CaCl_2_ (Sigma, C4901) in 0.1 M sodium cacodylate buffer (“caco buffer”, Sigma, C0250) for 48h (at RT for the first 24 h and at 4C for the remaining 24 h). Organoids were prepared for EM by treating them with 1% OsO_4_ (EMS, 19170) with 10 mg/ml potassium ferrocyanide (EMS, 20150) and 0.003M CaCl_2_ in 0.1M caco buffer for 1h at RT. Organoids were washed afterwards and stained with a 2% OsO_4_ solution with 0.003M CaCl_2_ in 0.1M caco buffer for 1h at RT. Sections were then stained with 2% uranyl acetate (“UA”, EMS, 8473) at 4°C overnight, dehydrated, and embedded in LX-112 epoxy resin (LADD Research Industries, 21310) at 60°C for 72h. Blocks were trimmed and sectioned at 40 nm slice thickness with a Leica EM UC6 Ultramicrotome and sections were collected on carbon-coated Kapton tape using an ATUM (Kasthuri et al., 2015). Strips of tape were mounted on silicon wafers and sections were post-stained with 2% UA in water and 3% lead citrate (Leica Biosystems ULTROSTAIN II). Sections were imaged using a FEI Magellan Scanning Electron Microscope (Thermo Fisher Scientific) equipped with a custom image acquisition software (WaferMapper(Hayworth et al., 2014)). A total of 12 panoramic high-resolution images (one per organoid) were acquired using the backscattered electron detector (7 kV, 1us/pixel dwell time, ranging from 86k to 106k pixels in the x axis and from 106k to 150k pixels in the y axis, at 4nm/resolution). Due to natural size variation among organoids, the final 2D panoramas varied slightly in dimensions to ensure spatial coverage from the surface to the central region of each organoid. Stitched EM images were imported into VAST(Berger et al., 2018) for visualization and manual annotation of synaptic clefts, creating a ground truth dataset. A deep neural network based on a U-Net architecture(Ronneberger et al., 2015) was then trained on this ground truth using the mEMbrain MATLAB package(Pavarino et al., 2023). The annotated dataset comprised less than 5% of the total volume and was produced by two human experts (∼20 hours total annotation time; pixel-level validation accuracy >0.95). mEMbrain’s interactive environment facilitated both model training and accuracy evaluation. Predicted synapses were represented as 2D probability maps registered to the EM images in VAST. Synapses were initially defined as connected components of pixels with a synapse probability greater than 40%, a threshold chosen to approximate expert annotations. Model performance exceeded 95% accuracy based on manual verification of the largest synapses (>95th percentile by area), providing an estimate of the false positive rate for unbiased condition comparisons. To standardize automatic synapse quantification across sections, a computational synapse-size threshold was determined for each section individually. Specifically, 103 randomly selected predicted synapses per section were manually classified as “synapse” or “non-synapse” by a human expert. A psychometric curve was then fitted to these annotations to establish a threshold corresponding to >80% model accuracy. Final synapse counts per section were obtained using these individualized thresholds, ensuring objectivity and consistency across samples. To account for structural variability and region-specific differences in synapse distribution, each section was divided into five horizontal segments. Synaptic density was calculated in each segment, and the one with the highest density was selected for statistical comparison (within each section, synaptic density variation across segments ranged from 0.00 to 0.14). This approach minimized sampling bias across organoid layers. The same set of 103 randomly selected predictions was further used to classify synaptic contacts by their postsynaptic target. Synapses were categorized as spine synapses if they were located on putative dendritic spines—morphologically consistent with spine-like structures observed in 2D EM—or on clearly defined spines attached to dendritic shafts. The remaining synapses were classified as targeting non-spine elements. Comparisons of synaptic densities across groups were performed using the Wilcoxon rank-sum test. The distribution of spine vs. non-spine synapses were analyzed using two-sided Fisher’s exact test. Statistical significance was defined as *P* < 0.05.

### Expansion microscopy analyses

We adapted a recent approach leveraging expansion microscopy and combinatorial antigen barcoding(Park et al., 2025). Seven-month-old organoids cultured in either CDM4 or APM were transduced with a mix of nine AAVs (8×10 vg each), each expressing a CAG-driven spaghetti-monster fluorescent protein(Viswanathan et al., 2015) tagged with distinct epitopes (V5, Myc, Flag, Ollas, E, S, HSV, Protein C, or Ty1). After three weeks, organoids were fixed in 4% PFA, embedded in gelatin, cryosectioned at 50 µm, and mounted on HistoBond slides. Sections were expanded using the Magnify protocol(Klimas et al., 2023b), followed by overnight gelation at 37°C in a humidified, nitrogen-purged chamber. Samples were homogenized, fully expanded via water washes, and re-embedded in a non-expandable hydrogel to allow iterative immunostaining. Expanded samples were blocked and stained with primary antibodies overnight, washed, and incubated with secondary antibodies the following night. Before imaging, samples were transferred to six-well glass-bottom plates. A Nikon W1-SORA spinning-disk confocal microscope was used: 4x overview images identified dense regions, followed by 10x z-stacks and 40x tile scans. Large 40x scans were stitched using BigStitcher; registration across rounds was performed with BigStream using a two-stage alignment pipeline (coarse RANSAC-based feature matching, followed by fine intensity-based registration using Mutual Maximum Information). Affine transformation matrices were applied to full-resolution datasets. Preprocessing for low SNR rounds included percentile-based contrast enhancement and 3D median filtering. Aligned volumes were imported into Webknossos, converted to wkw format, and merged into a single nine-channel dataset. Ten neurons per condition (CDM4, APM) were manually skeletonized based on multi-tag expression in the cytosol. Resulting traces (30 per condition) were exported as .swc files, and morphometric parameters were quantified using Navis in Python.

### Dissociation of brain organoids and scRNA-seq

Organoids were dissociated as previously described(Antón-Bolaños et al., 2024). Dissociated cells were resuspended in 100 μl of PBS, filtered with a 35 μm cell strainer tube (Corning) to avoid aggregates, and counted with a Countess II automated hematocytometer (Thermo Fisher Scientific) or a Cellometer K2 (Nexcelom Bioscience) and AOPI stain (Nexcelom Bioscience, CS2-0106-5ML). Single-cell suspensions were loaded into the Chromium Next GEM Chip G (10x Genomics, 1000120) and run with the Chromium Controller to generate single-cell GEMs. scRNA-seq libraries were prepared with the Chromium Single Cell 3’ Library and Gel Bead Kit v3.1 (10x Genomics, 1000268). The resulting libraries were pooled based on molar concentration and sequenced on a NovaSeq X or NovaSeq 6000 instrument (Illumina) either 28 bases for read 1, 55 bases for read 2 and 8 bases for index read 1, or 28 bases for read 1, 90 bases for read 2, and 10 bases for index read 1. If necessary, after the first round of sequencing, we re-pooled libraries based on the actual number of cells in each library, and re-sequenced with the goal of producing an equal number of reads per cell for each sample (with a target read depth of 20,000 reads/cell).

### scRNA-seq data analysis

The preprocessing of all scRNA-seq samples, including the conversion of raw BCL files into FASTQ files, reads alignment, and the generation of gene-barcode matrices, was performed using 10x Genomics Cell Ranger suite tool(Zheng et al., 2017) following the steps described in our previous publication(Antón-Bolaños et al., 2024). For *de novo* generated samples with moderate to high ambient RNA levels, CellBender(Fleming et al., 2023) v0.3.0 was used to remove systematic biases and background noises. The remove-background pipeline with default parameters was used by taking the raw gene-by-cell count matrices produced by Cell Ranger pipelines as inputs. The filtered gene-barcode count matrices by Cell Ranger and CellBender were imported into Seurat v4.3.0(Hao et al., 2021) for downstream analysis.

#### Genetic demultiplexing

In the cases where multiple samples of different genetic backgrounds were pooled together and sequenced in the same lane to maximize the sequencing capacity, Demuxlet v1.0(Kang et al., 2018) was applied with default parameters to determine the sample identity for each barcoded droplet and identify ambiguous droplets and doublets that contain two heterogeneous cells. Ambiguous droplets and doublets identified were removed from downstream analysis. To run Demuxlet, the BAM file output from Cell Ranger, the filtered barcode list generated by the initial preprocessing pipeline, and reference variant-call-format (VCF) files for each expected cell line were used as inputs. The reference VCF files were prepared by the method described in our previous publication(Antón-Bolaños et al., 2024).

To address doublet overestimation when applying Demuxlet to multiplexed samples with ambient RNAs, following the usage of CellBender, Souporcell v.2.5(Heaton et al., 2020) was suggested by the developer and used with default parameters as an alternative method, which allows genotype-free demultiplexing and clusters cells by the genetic variants detected within the input reads. The same set of input data plus the GRCh38 human reference genome FASTA file were used. We followed recommendations to include the reference VCF that has genetic variants of pooled individuals to enhance accuracy. Lastly, the Assign_Indiv_by_Geno.R function from the Demuxafy(Neavin et al., 2024) framework was used to correlate clusters with donors in the reference SNP genotypes.

#### scRNA-seq data quality control, normalization, dimensionality reduction, and clustering

For each scRNA-seq dataset, we performed initial QC filtering and retained cell profiles having total UMI counts greater than 500 and less than 20000, unique feature counts greater than 200, and the percentage of mitochondrial RNA less than 15. Seurat’s SCTransform function was used to normalize the data and regress out the mitochondrial RNA percentage. The RunPCA function was used to perform principal components analysis (PCA), and the top 30 principal components were used by the FindNeighbors function to construct a k-nearest-neighbor (KNN) graph with default parameters. Cells were then clustered by the Louvain algorithm using the FindClusters function with resolution = 0.3 for a rough granularity to start with. Clusters showing remarkably low UMI counts, low unique feature counts, and high percentage of mitochondrial RNA were labeled as low-quality cells and removed.

#### Data compilation of group-published data

To unify data processing across datasets, raw scRNA-seq FASTQ files for organoids from Uzquiano et al(Uzquiano et al., 2022). and Velasco et al.(Velasco et al., 2019) were reprocessed using Cell Ranger 7.2.0 (the most notable change from the original processing is the inclusion of intronic reads in the count matrices).

#### Cluster annotation and data integration

Before any data merging/data integration, for each scRNA-seq data, we manually annotated each cluster by an ensemble approach, as described in our previous publication(Antón-Bolaños et al., 2024). Doing so provides a solid way for cell type assignments by taking the prior domain knowledge, Seurat’s FindMarkers output, and alignment with notations in previously published reference maps(Uzquiano et al., 2022) into consideration.

In pursuit of comparing organoids cultured under the conventional CDM4 and the novel APM conditions, we profiled APM-treated organoids at 4 months, 6 months, 9 months, and 1 year in this study. To facilitate downstream comparative analysis and enable more interpretable results, we refined cell type annotations between treatment conditions and corrected for sequencing batches as follows: For each age, scRNA-seq datasets from two conditions were merged by using Seurat’s merge function. Considering the notable data imbalance between CDM4- and APM-treated organoids at 4 months and to control the overall cell number disparity between conditions, a 50% downsampling was applied on the CDM4 scRNA-seq dataset while retaining the original sample proportions before merging. Features that are repeatedly variable across datasets in the list were obtained by using Seurat’s SelectIntegrationFeatures with default parameters and set as the variable features of the merged object. The RunPCA function was applied, and the merged data was then batch-corrected by Harmony(Korsunsky et al., 2019) 1.2.0. The downstream clustering used the standard Seurat pipeline by taking Harmony-corrected embeddings with a resolution set between 0.8 and 1.0. To examine top up-regulated genes within each cluster for cell type annotation, before using FindAllMarkers, Seurat’s PrepSCTFindMarkers was used to recalculate the SCT assay by taking the minimum median UMI across all datasets being merged as a scale factor to correct for sequencing depth variations across different datasets. To achieve higher resolution and finer granularity in certain clusters, including clusters that were labeled as cycling cells and clusters with underlying heterogeneity of cell types, subclustering was performed. Due to the absence of organoid samples with matched genetic backgrounds across conditions, the integrated 4-month data were excluded from the age-wise comparative analysis across conditions. Thereby, we retained cell profiles from organoids with matched genetic backgrounds for other time points for data representation and downstream analysis as follows: 6 months: (CDM4: n = 10, 64,817 cells; APM: n = 9, 41,231 cells), 9 months (CDM4: n = 6, 18,104 cells; APM: n = 3, 8533 cells), 12 months (CDM4: n = 4, 7802 cells; APM: n = 3, 6084 cells).

To assess the reproducibility of organoids, we calculated Aitchison distances based on cell-type compositional data between every unique pair of samples at each time point by using the aDist function from the R package robCompositions v2.3.1(Templ et al., 2011). Aitchison distances between replicates within the treatment protocol were compared by Wilcoxon rank-sum tests. Three previously published human fetal cortex datasets(Polioudakis et al., 2019; Trevino et al., 2021; Uzquiano et al., 2022), as noted in our previous publication(Antón-Bolaños et al., 2024), were used to provide referential median sample-to-sample variabilities by their respective cell type annotations.

To get a comprehensive overview of the organoid development across time points, We generated a developmental time map by merging the annotated, batch-corrected scRNA-seq data of CDM4-treated organoids, including 257,044 cells from 77 organoid samples across 3 months (n = 31; 69,719 cells), 4 months (n = 6; 33,825 cells), 5 months (n = 6; 32,115 cells), 6 months (n = 12; 74,172 cells), 9 months (n = 10; 32,550 cells), 12 months (n = 4; 7802 cells), 18 months (n = 2; 1415 cells), 24 months (n = 2; 2274 cells), 36 months (n = 2; 2481 cells), and 42 months (n = 2; 691 cells). In general, we annotated 17 cell types and we grouped cell types into several populations for broader representation as follows: 1) Progenitors (aRG: apical radial glia; oRG: outer radial glia; IP: intermediate progenitors; IN progenitors: interneuron progenitors; tRG: truncated radial glia); 2) ExNeu (CFuPN: corticofugal projection neurons; CPN: callosal projection neurons; PN: unspecified projection neurons); 3) IN: immature interneurons; 4) GlialProg: Glial precursors; 5) AST: astrocytes; 6) OPC: oligodendrocyte progenitors. The merged data was jointly renormalized by SCTransform and reprocessed by the standard Seurat pipeline. The time-series object of APM-treated organoids from 4 months to 12 months was generated by the same method after merging the APM counterpart from age-wise integrated data as described above.

#### Analysis of previously published human fetal data

A previously published perinatal single-nucleus RNA-seq (snRNA-seq) dataset for human cortical development was used for comparison. The dataset from Velmeshev et al.(Velmeshev et al., 2023) includes profiles from 709,372 nuclei of 169 brain tissue samples from 106 individuals, with a wide age range spanning from the second trimester of gestation to adulthood. The snRNA-seq data and metadata were downloaded from https://cells.ucsc.edu/?ds=pre-postnatal-cortex. We subsetted the snRNA-seq data to an age range from the second trimester up to 4 years which is a proxy of time window in our time-series data. The data was further subset by brain regions including cortex (CTX) and frontal cortex (FC) with a spectrum of cell type populations including Progenitors, ExNeu (excitatory neurons), IN (interneurons), GlialProg (glial progenitors), AST (Astrocytes), and OPC (oligodendrocyte precursor cells). The subset data was renormalized by SCTransform and reclustered by the standard Seurat pipeline. To compare with the reference human perinatal snRNA-seq dataset and explore what age the time-series organoids data is mapping to, we used labels by concatenating the information of cell type classes and age and performed reference-based label transfer using Seurat’s FindTransferAnchors and MapQuery functions. To refine the label transfer results for query cells predicted to be in the second trimester, we performed a secondary label transfer by subsetting these cells and using an additional scRNA-seq dataset from human fetal cortical samples at post-conceptional week (PCW) 16, 20, 21, and 24 as the reference(Trevino et al., 2021).

#### Identification of multicellular programs (MCPs) associated with aging

To explore the maturation profiles of the time-series organoids data, we used the aforementioned human perinatal dataset as inputs and applied DIALOGUE v1.0(Jerby-Arnon and Regev, 2022), a tool designed to uncover multicellular programs (MCPs) of co-regulated genes across different cell types and map how the transcriptome varies with changes in its environment. Following DIALOGUE’s vignette, a list of cell type objects was generated by the make.cell.type function to represent major cell populations in the human perinatal dataset by using a series of input arguments including the single-cell gene expression data, sample identity, and PCA embeddings for each cell type of interest. Metadata of the SCTransform-normalized total UMI counts and the developmental age in days were included as a technical variability and a meaningful biological covariate to adjust for the analysis. Next, the DIALOGUE.run function was applied to the above list. As a result, five MCPs were identified, and each MCP output cell-type specific up- and down-regulated gene sets. Of the five MCPs, MCP4 more consistently showed the highest correlation with age within each cell population. Subsequently, we combined up- and down-regulated genes identified in each population within MCP4, respectively, into two master gene sets. After applying Seurat’s AddModuleScore function using the two master gene sets, we subtracted the module scores of the up-regulated gene set from those of the down-regulated gene sets. Surprisingly, the ascending trend of module scores still holds a high correlation with age. Therefore, we referred to the subtracted scores as the “Maturation” scores and we adopted the same approach to calculate the “Maturation” scores by using the above master gene sets in the context of the time-series organoids scRNA-seq data.

#### Analysis of presynapse, synapse, and postsynapse signature modules

Gene sets of gene ontology terms: synapse (GO:0045202), presynapse (GO:0098793), and postsynapse (GO:0098794) containing 1778, 879, and 1181 genes, respectively, were retrieved from the SynGO knowledge base of release version 1.2 (https://www.syngoportal.org/)(Koopmans et al., 2019). Each gene set was used to calculate module scores for cells in integrated scRNA-seq data of CDM4- and APM-treated organoids at 6 months, 9 months, and 12 months by using Seurat’s AddModuleScore function. To compare, under each age, if organoids under two treatment conditions displayed different averaged expression levels for each program, linear mixed-effects models were built by the R package lme4(Bates et al., 2015) to model the scores by including which organoid the cell belongs to and the genetic background of that organoid as random effects (Score ∼ (1|Organoid) + (1|Genotype)). To adjust for cell count differences in each broad cell population across organoids, the reciprocal of the square root of the observed cell count per population per organoid was included as a weight factor. Models with or without treatment as an additive fixed effect were compared by the anova function from the R stats package and *P* values across tests of the above 3 signature modules were adjusted by the Benjamini-Hochberg method.

#### Comparing cell type proportions between treatment conditions

The cell type proportion changes between CDM4- and APM-treated organoids at 6 months, 9 months, and 12 months were evaluated by negative binomial mixed-effects models. A matrix of cell counts for each annotated cell type per organoid was generated. Given a cell type of interest, a negative binomial mixed-effects model was created by using the glmer.nb function from lme4(Bates et al., 2015), which takes the genetic background of an organoid replicate as a random effect to account for the variability in genotypes. The model also includes the logarithm of the total cell number of each organoid sample as an offset to account for the cell count differences (model formula: Cell Type ∼ offset(log(LibrarySize)) + (1|Genotype)). The significance of cell type proportion differences between organoids under two treatment conditions was calculated by comparing negative binomial models with and without treatment as an additive fixed effect using the anova function. Benjamini-Hochberg (BH) method was used to adjust *P* values from multiple independent tests. Cell type proportion differences between MonChr chimeroids and 9mo organoids were calculated using the same methods as above, and only considering cell types with a minimum abundance of 5% in 9mo organoids.

#### Differential gene expression analysis and gene ontology analysis

To identify differentially expressed genes (DEGs) between CDM4- and APM-treated organoids at each time point (6, 9, and 12 months), we employed a pseudo-bulk approach to analyze the data at the sample level. We took pan-ExNeu, AST, and IN as partitions of interest by using the above-mentioned grouping criteria. Pseudobulk profiles were generated on a partition basis by aggregating UMI counts for each gene across cells of that partition within each organoid sample. We require at least 20 cells within a partitioned cluster per organoid to retain the sample. Genes with fewer than 10 total UMI counts in at least 2 samples were excluded. The pseudo-bulk differential expression analysis was performed by using DESeq2(Love et al., 2014) with the design formula ∼ Genotype + Treatment to identify DEGs between treatment conditions while regressing out the variation by genetic backgrounds. DEGs were defined by a cutoff with the absolute value of Log2 Fold Change > 0.5 and FDR-adjusted *P*-value < 0.05. Up-regulated DEGs were used by the enrichGO function from clusterProfiler(Yu et al., 2012) to perform GO enrichment analysis. The simplify function with a default similarity cutoff of 0.7 from the same package was used to remove the semantic redundancy of enriched GO terms.

#### Comparative analysis of DIALOGUE-determined maturation scores

To explore if APM-treated organoids show different DIALOGUE maturation scores in a time-series or time-specific manner, the age-wise integrated data from 4 months to 1 year were merged and jointly renormalized by SCTransform and reprocessed by the standard Seurat pipeline. We used Seurat’s AddModuleScore function to calculate module scores using DIALOGUE MCP4’s full list of “Up” and “Down” genes, separately, and then took the subtraction of “Down - Up” as a single “Maturation” score, as described above. For the comparative analysis, the Z-score, with a mean of 0, was used to represent the magnitude. Maturation module scores across the entire dataset were standardized using the R function scale. Focusing on the excitatory neurons, we first assessed the treatment effect over time by treating age as a continuous variable. The Z-scores were averaged across all cells within the partition of interest per organoid. A baseline linear model was created to evaluate how maturation scores change with age (score ∼ age). A second linear model incorporating treatment in addition (score ∼ treatment + age) was used to compare with the baseline model by anova function from the R stats package to test if adding treatment as an additive effect significantly improves the model fitting. Next, to assess in a time-specific manner, linear mixed-effects models were used to model the scores by including which organoid the cell belongs to and the genetic background of that organoid as random effects (score ∼ (1|Organoid) + (1|Genotype)) and a weight factor to adjust for cell count differences in excitatory neurons across organoids. Models with or without treatment as an additive fixed effect were evaluated by anova function. The above statistical models were built by the R lme4(Bates et al., 2015) package.

## Supplementary Figures

**Supplementary Figure 1.**
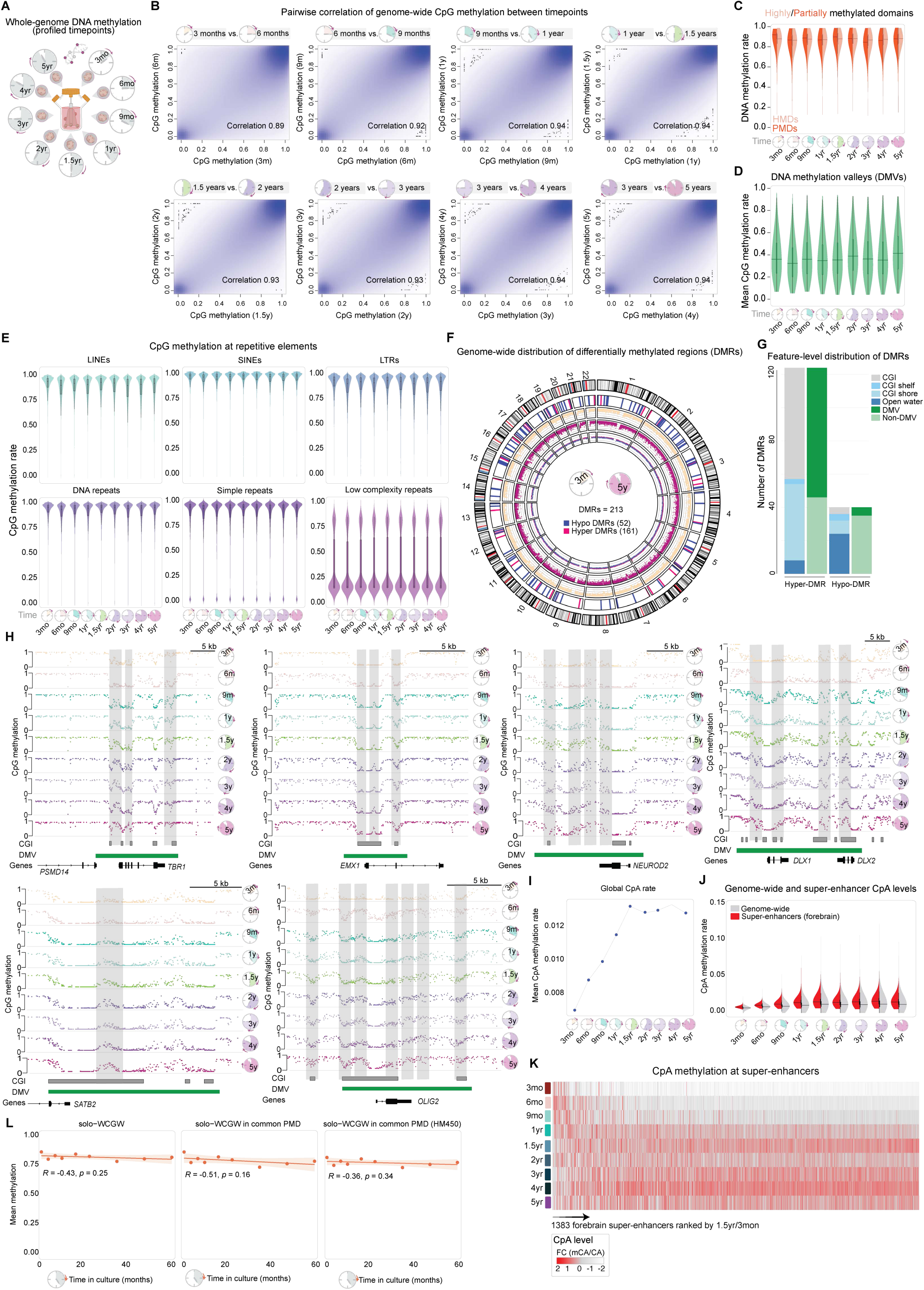
**A**, Overview of time points profiled by whole-genome bisulfite sequencing (WGBS). **B**, Pairwise density plots comparing genome-wide CpG methylation values between consecutive time points. Pearson correlation coefficients are indicated. Each dot is a single CpG. **C** DNA methylation rates in highly and partially (HMD, PMD) methylated domains are shown. The line in the violin plot indicates the median of the distribution. **D,** Distribution of average methylation levels within DMVs across time points. The line in the violin plot indicates the median of the distribution. **E**, Distribution of average methylation levels across different repeat classes (LINEs, SINEs, LTRs, DNA repeats, low-complexity repeats). **F**, Circos plot showing genomic positions of n=213 DMRs identified between 3-month and 5-year organoids (first inner track; red: hypermethylated, blue: hypomethylated). Outermost track shows the hg19 chromosome ideogram. Inner scatter tracks show genome-wide CpG methylation averaged in 50kb bins for 3-month (second track) and 5-year (third track) organoids, followed by the difference in methylation (3m - 5y; innermost track). **G**, Genomic annotation of age-correlated DMRs, showing enrichment across CpG islands (CGIs) and DMVs. **H**, Genome browser tracks showing methylation profiles of DMRs at the TBR1, EMX1, NEUROD2, DLX1/2, SATB2, and OLIG2 loci across all time points. DMR regions are highlighted (grey boxes). CGIs, DMVs, and genes are annotated below. **I**, Average global CpA methylation rates across all time points. **J**, Distribution of average CpA methylation rates within forebrain super-enhancers (1383 super-enhancers) and genome-wide 30kb background bins. The line in the violin plot indicates the median of the distribution. **K**, Heatmap of the fold change of CpA methylation (1.5 yr/3 mon) at the forebrain super-enhancers. Column shows individual super-enhancers. **L**, Scatterplot comparing time-in-culture (in months) of sequenced organoids and average methylation rates at solo-WCGWs (left), solo-WCGWs in common PMDs (middle), and solo-WCGWs within common PMDs represented on the HM450 array (right). Red line: linear regression fit; R: Pearson correlation coefficient; p: correlation test p-value.

**Supplementary Figure 2.**
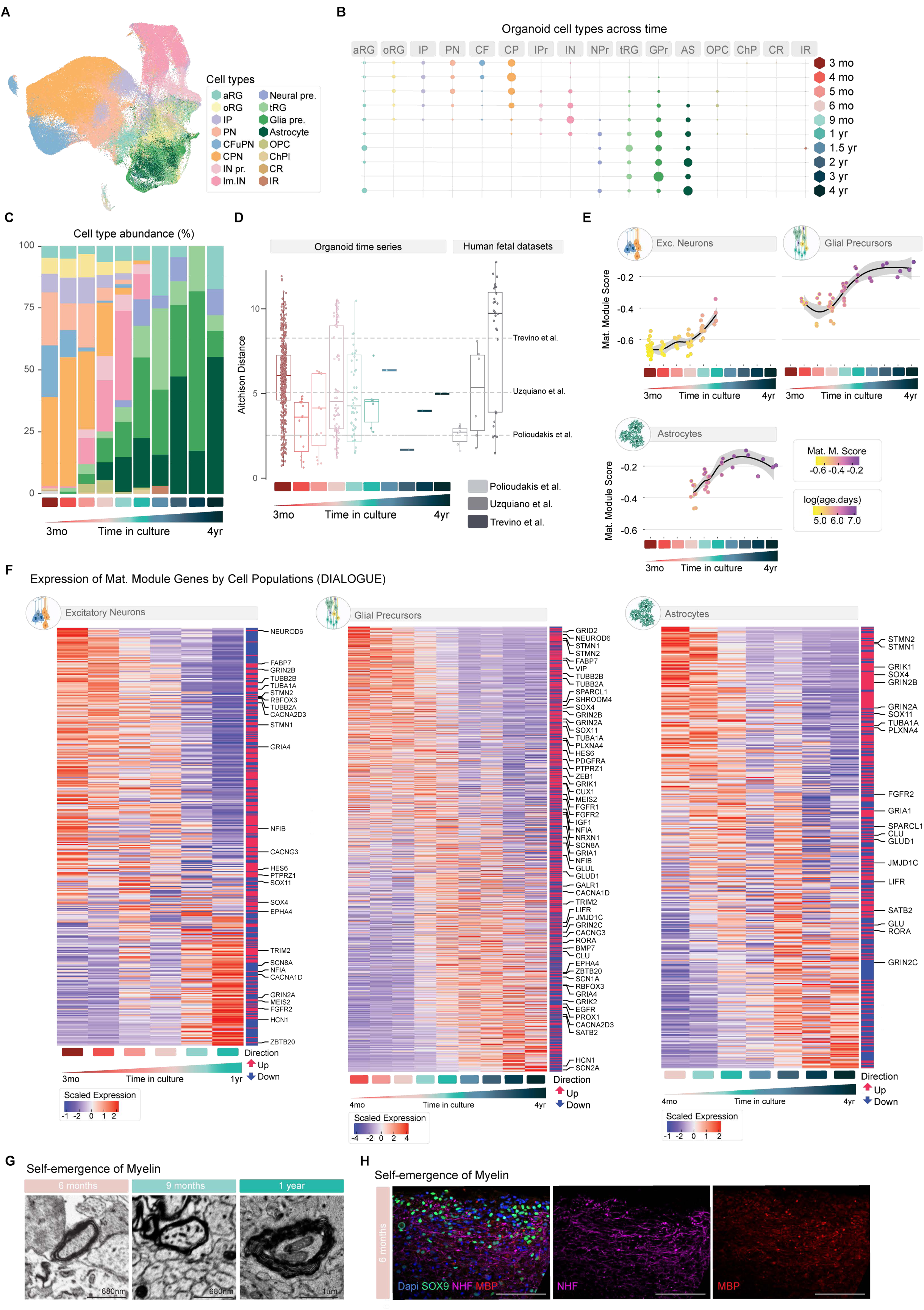
**A**, UMAP plots colored by annotated cell type. **B**, Dotplot showing the distribution of cell types within each timepoint. Dot size represents percent abundance, y-axis represents timepoint, x-axis and color represent cell type. **C**, Stacked barplots showing the summarized abundance of cell types for each timepoint. **D**, Boxplots showing the Aitchison distance between each pair of organoids within each timepoint, and between pairs of endogenous human tissue samples from published datasets. The Aitchison distance indicates how dissimilar the relative distributions of cell types are between two samples. **E**, Scatterplots showing the mean Maturation Score (y-axis) of all astrocytes, all excitatory neurons, and all glial precursors within each organoid, grouped by timepoint (x-axis). Smoothed conditional means (black lines) with default 95% confidence intervals (grey regions) were calculated by the geom_smooth() function in R. Points were given slight x-axis jitters to improve visibility; horizontal position within a timepoint does not indicate differences in age. **F**, Heatmaps showing the expression of all genes contributing to DIALOGUE’s MCP4 within a given cell population in organoids across time. Values for each timepoint represent the mean of the counts-per-million of the pseudobulked count matrices for that cell type within each organoid within that timepoint, Z-scored for each gene. Genes are ordered by the weighted expression over time; genes expressed mostly at earlier time points are at the top, and genes expressed mostly at later time points are at the bottom. The color bar on the right indicates whether the gene was marked as in the “up” (red) or “down” (blue) aspect of MCP4. Labeled genes were manually selected to show important biological features over time. Because DIALOGUE did not derive specific genes for MCP4 within glial precursors or progenitors, the heatmaps for these cell populations show all genes belonging to MCP4 in any cell population. **G**, Electron microscopy representative images showing the presence of myelin along the longitudinal trajectory. **H,** Immunohistochemistry of organoids at 6 months showing the presence of myelinated axons. Scale bar: 100 μm.

**Supplementary Figure 3.**
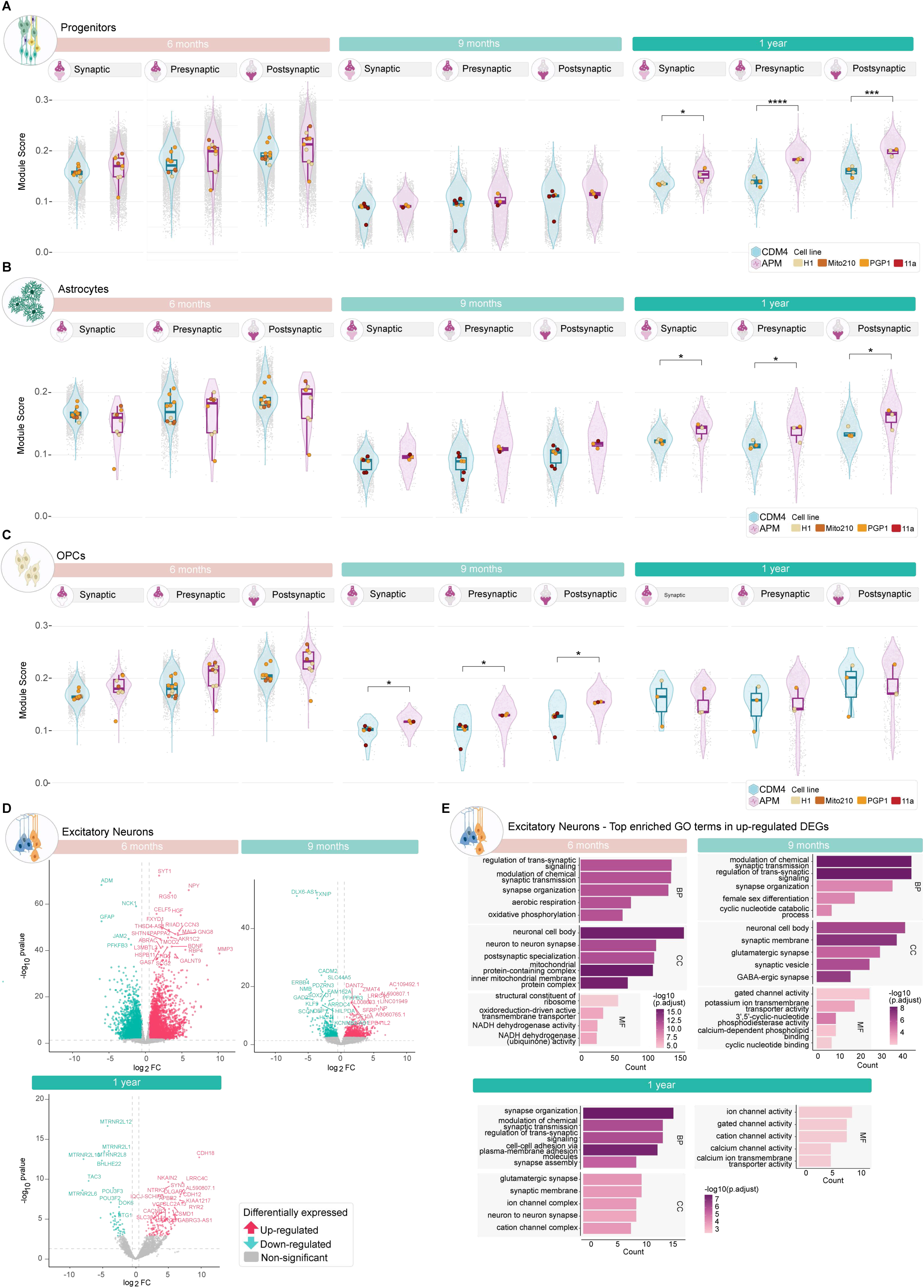
A-C. Expression of three SynGO synaptic genes sets in other cell populations from CDM4- and APM-treated organoids at distinct ages including 6 (CDM4: n = 10 organoids; APM: n = 9 organoids), 9 (CDM4: n = 6 organoids; APM: n = 3 organoids), and 12 months (CDM4: n = 4 organoids; APM: n = 3 organoids). Violin plots show the module score distribution for synapse (GO:0045202), presynapse (GO:0098793), and postsynapse (GO:0098794) gene sets across a given cell population. Mean module scores for individual organoids are shown as points, colored by genetic backgrounds. The bottom and top edges of the boxes represent the 25th and 75th percentiles, with the middle line representing the median and whiskers spreading the 1.5 x interquartile range (IQR) from the hinges. Module scores were calculated by using Seurat’s AddModuleScore function. For each age, FDR-adjusted *P* values that show significant differences between each population in CDM4- and APM-treated organoids are based on linear mixed-effects models and are indicated by asterisks: *P* < 0.05 (*); *P* < 0.01 (**); *P* < 0.001 (***); *P* < 0.0001 (****) (see Methods.) **D**, Pseudobulked differential expression analysis of comparing excitatory neurons between CDM4- and APM-treated organoids at distinct ages, including 6, 9, and 12 months. Volcano plots display the differentially expressed genes (DEGs) defined by the significance thresholds: |Log2 Fold Change|> 0.5 and FDR-adjusted *P*-value < 0.05 (denoted by gray horizontal/vertical dashed lines). The top 30 DEGs with the highest significance are labeled. **E**, Top 5 enriched Gene Ontology (GO) terms per ontology category of up-regulated DEGs in excitatory neurons of APM-treated organoids at distinct ages, including 6, 9, and 12 months. Gene ontology enrichment analysis was performed by using the enrichGO function from clusterProfiler with settings that include pvalueCutoff = 0.01, qvalueCutoff = 0.05, and pAdjustMethod = “BH”. The simplify function with a default similarity cutoff of 0.7 from the same package was used to remove the semantic redundancy of enriched GO terms.

**Supplementary Figure 4.**
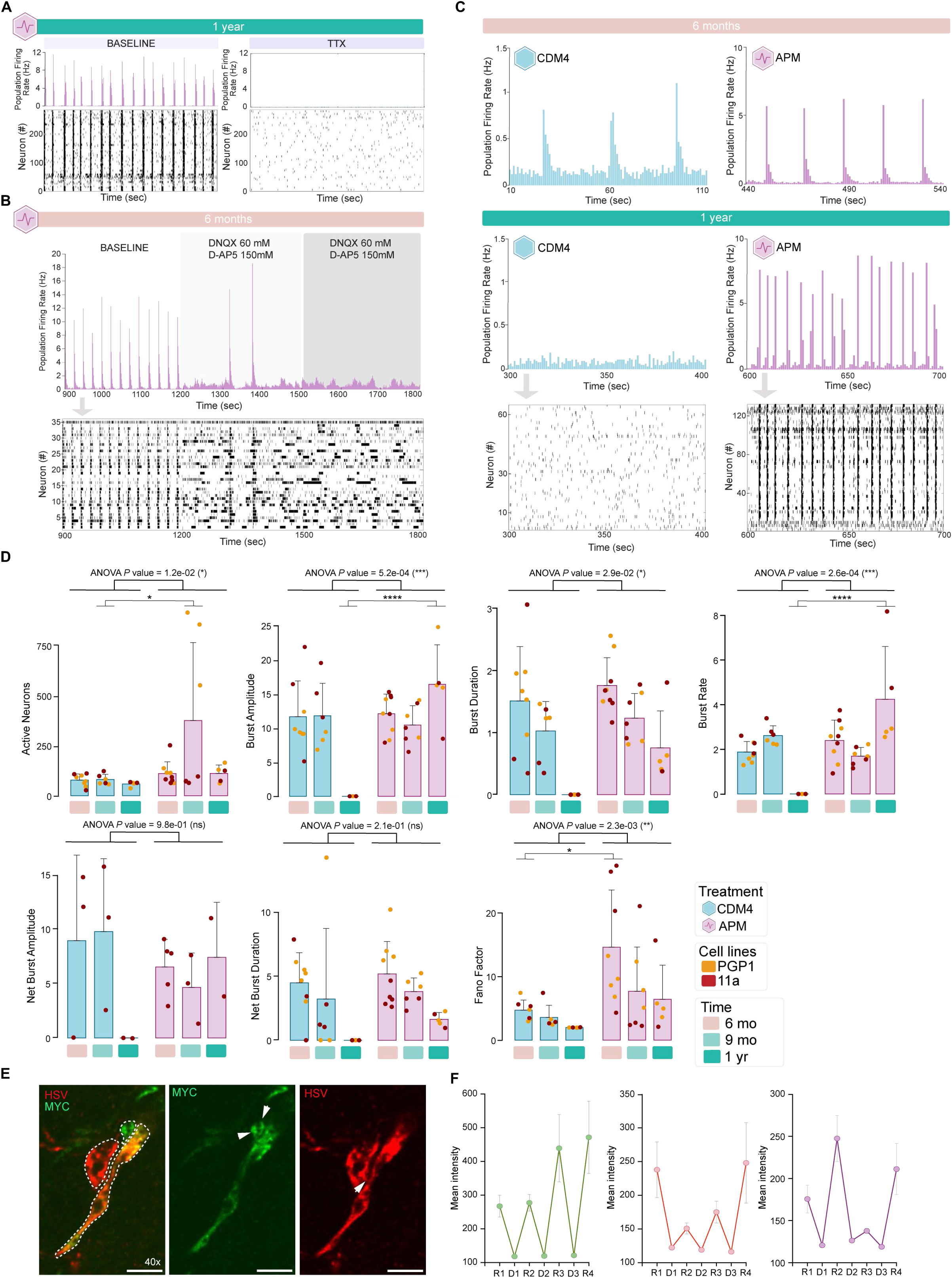
**A**, Tetrodotoxin (TTX) abolished the spiking activity. Example recording is presented through the population-averaged firing rate (top) and a spike raster plot from all detected units (bottom). **B**, Network bursts are mediated by glutamatergic neurotransmission. Population firing rate and spike raster plot of a representative organoid treated with glutamate receptor antagonists (DNQX and D-AP5). **C**, Representative population-averaged firing rate and spike raster plot for a subset of units in 6-month (top) and 12-month (bottom) organoids, showing the difference between CDM4 (left) and APM (right) groups. **D,** Characterization of the network bursting activity for CDM4- and APM-treated organoids. For each measured metric by MEA, a Type III two-way ANOVA test was performed to assess the effects of Treatment, Age, and their interaction Treatment:Age. For cases where quantifications of more than one cell line were available, the genetic background was included as a fixed effect. ANOVA *P* value is shown to denote the significance of the Treatment effect on the measured metric, as represented by asterisks. Post-hoc Tukey tests were performed to determine at which age the treatment significantly differ and adjusted *P* values by Tukey’s HSD method were represented by asterisks. **E**, Expression of different cytosolic epitope tag combinations enables tracing of densely packed neuronal processes, visualized here with two representative epitope tags. Three processes expressing either Myc tag, HSV tag, or both can be seen (indicated by dashed white lines). While expression of Myc tag or HSV tag alone would not allow unambiguous separation of these processes (see white arrows), combinatorial epitope tag expression enables their identification as three distinct neuronal processes. Scale bar: 10μm. **F**, Quantification of antibody de-staining efficiency in the three imaged color channels. R1-4 = staining rounds. D1-4 = de-staining rounds.

**Supplementary Figure 5.**
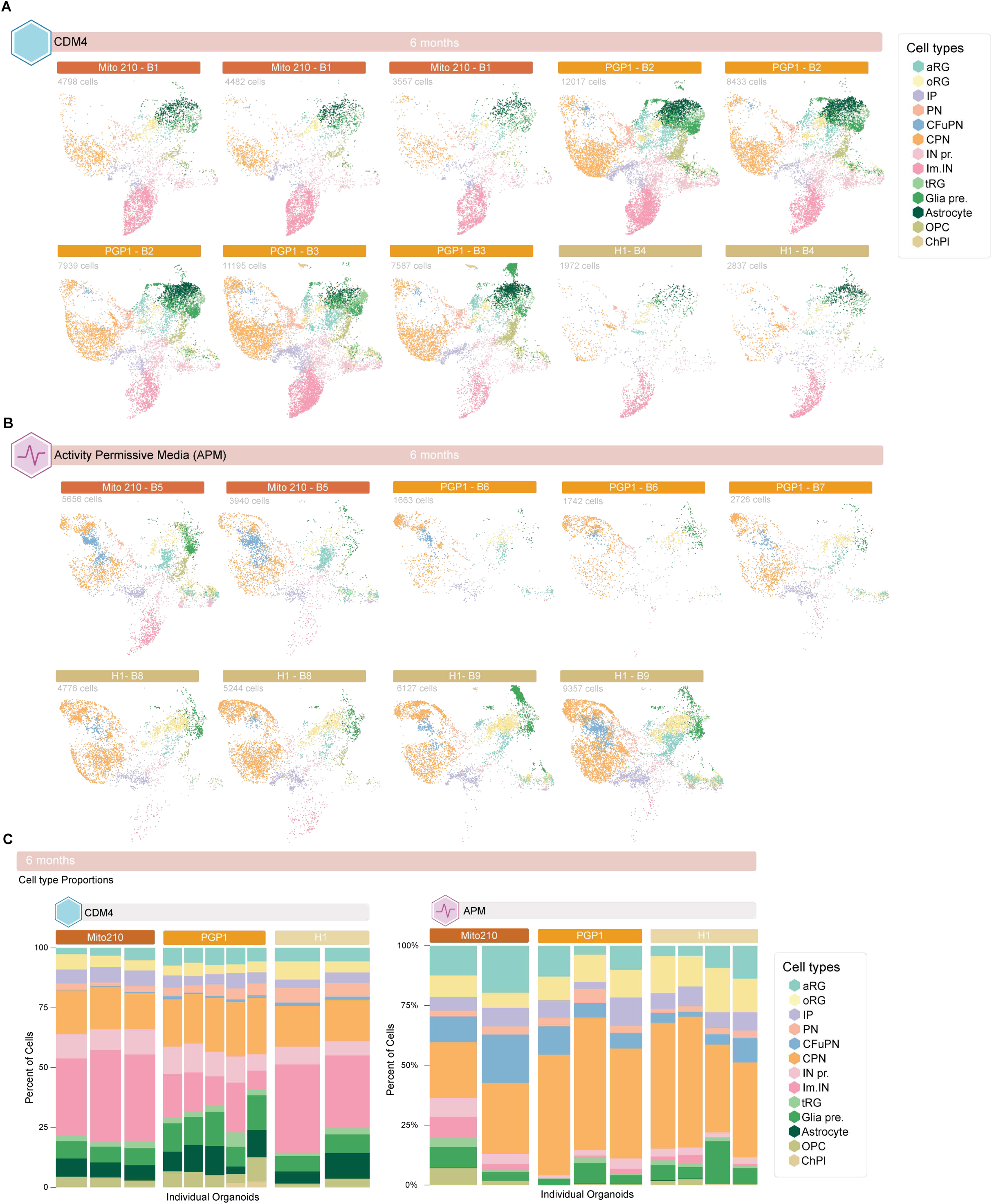
**A**, UMAP of 6-month CDM4-treated organoids, split by organoid and colored by cell types. **B**, UMAP of 6-month APM-treated organoids, split by organoid and colored by cell types. **C**, Stacked barplots showing the cell type compositions of each 6-month CDM4- and APM-treated organoid, subgrouped by genetic background/cell line.

**Supplementary Figure 6.**
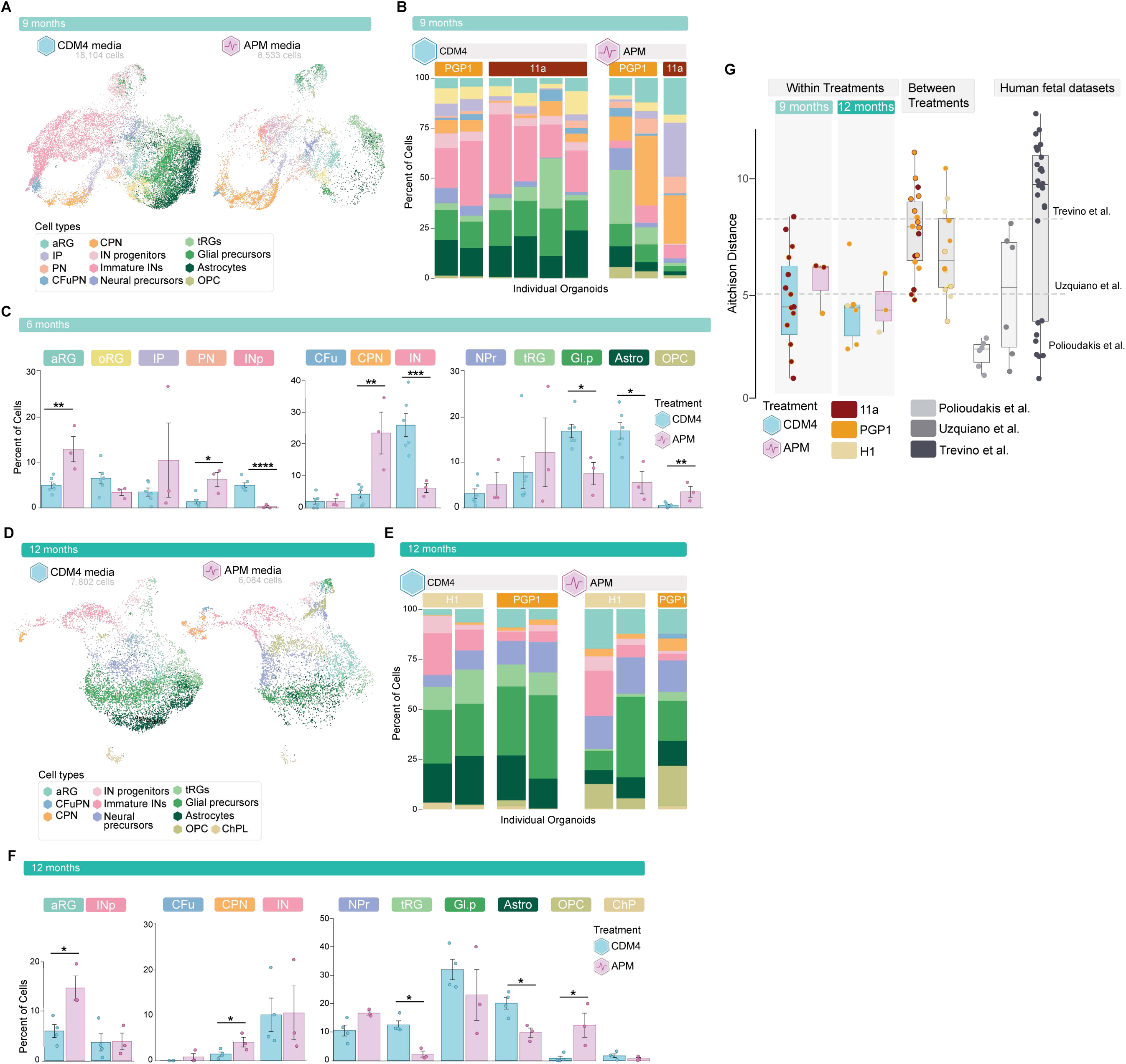
**A**, UMAP of integrated scRNA-seq data from 9-month CDM4- (n = 6 organoids, 18,104 cells) and APM-treated (n = 3 organoids, 8533 cells) organoids, colored by cell types. **B**, Stacked barplots showing the cell type compositions of each 9-month organoid, grouped by treatment and subgrouped by genetic background/cell line. **C**, Changes of cell type proportions between CDM4- (n = 6) and APM-treated (n = 3) organoids at 9 months. Bars represent the mean percentage of cell types for each treatment with error bars indicating the standard error of the mean. The significance of cell type proportion differences was calculated by the Likelihood Ratio Test comparing negative binomial models with and without treatment as a fixed effect using anova function. Each model takes the log of the total number of cells per organoid as an offset, the genetic background of the organoid as a random effect. FDR-adjusted *P*-values are indicated by asterisks: *P* < 0.05 (*); *P* < 0.01 (**); *P* < 0.001 (***); *P* < 0.0001 (****). **D**, UMAP of integrated scRNA-seq data from 12-month CDM4- (n = 4 organoids, 7802 cells) and APM-treated (n = 3 organoids, 6084 cells) organoids, colored by cell types. **E**, Stacked barplots showing the cell type compositions of each 12-month organoid, grouped by treatment and subgrouped by genetic background/cell line. **F**, Changes of cell type proportions between CDM4- (n = 4) and APM-treated (n = 3) organoids at 12 months. Bars represent the mean percentage of cell types for each treatment with error bars indicating the standard error of the mean. The significance of cell type proportion differences was calculated by the Likelihood Ratio Test comparing negative binomial models with and without treatment as a fixed effect using anova function. Each model takes the log of the total number of cells per organoid as an offset, the genetic background of the organoid as a random effect. FDR-adjusted *P*-values are indicated by asterisks: *P* < 0.05 (*); *P* < 0.01 (**); *P* < 0.001 (***); *P* < 0.0001 (****). **G**, Aitchison distance measuring the differences in cell type compositions was calculated for unique pairs of samples within each treatment and between 2 treatment conditions at 9 and 12 months, respectively. For each age, Aitchison distances between replicates within each treatment were compared by the Wilcoxon rank-sum test, and no significant dis-similarity was shown. The means for Aitchison distances (dotted lines) for three different datasets of endogenous human fetal cortex were included.

**Supplementary Figure 7.**
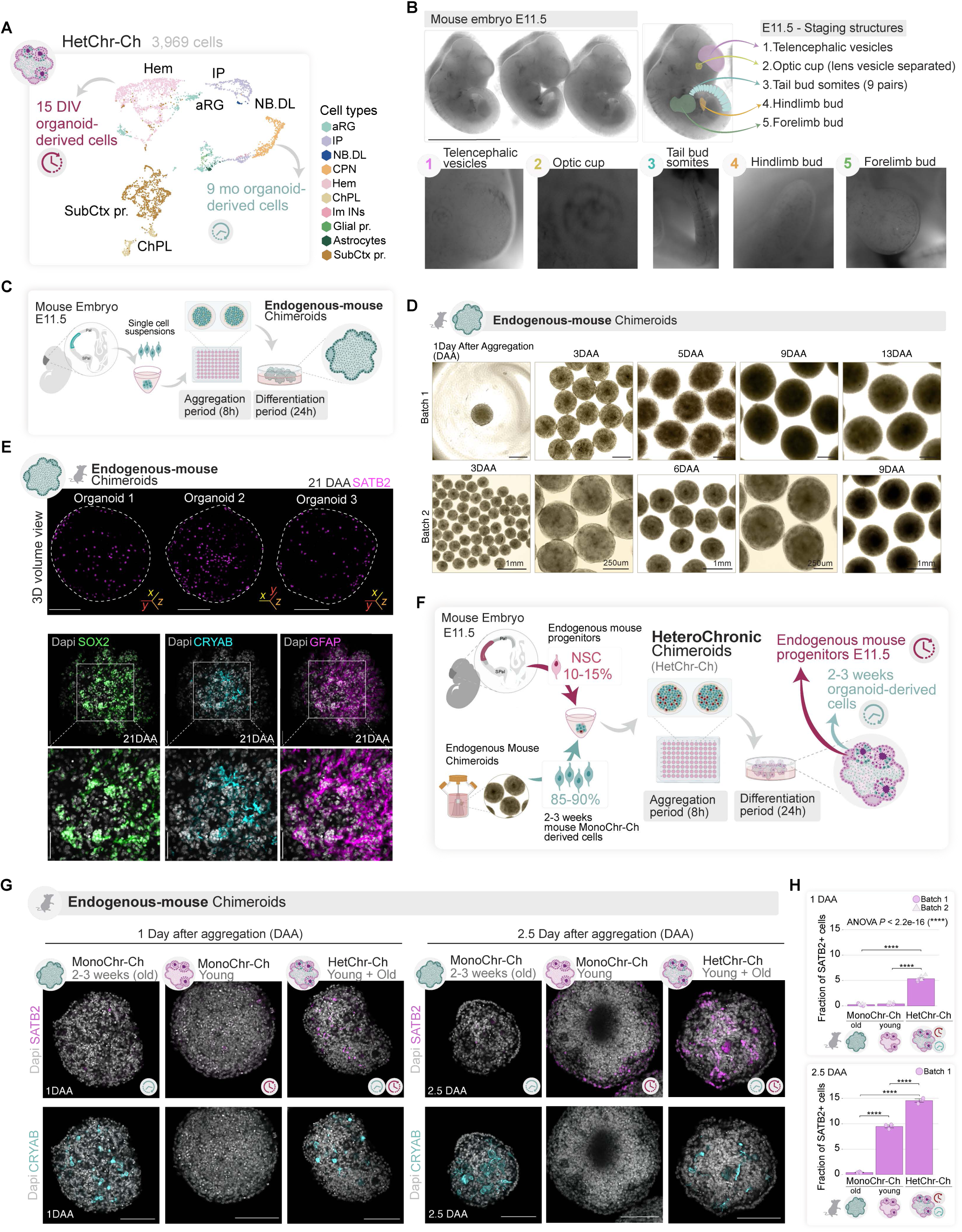
**A**, UMAP plot showing cells from a HetChr-Ch sample (n=3,969 cells), colored by cell type annotation of the old and young counterparts. **B**, Brightfield images of staged E11.5 embryos with annotated structures used for staging. Scale bar is 1mm. **C**, schematics of ex-vivo organoids derived from mouse embryos collected at E11.5. **D**, Brightfield images of ex vivo organoids after dissociation at different time points in vitro. DAA, days after aggregation. Scale bar is 100μm (top) and 1mm (bottom-zoom out), and 250μm (bottom-zoom in). **E**, Whole-mount immunofluorescence of ex vivo organoids 2-3 weeks after aggregation. Top panels show 3D volume view of individual organoids with SATB2. Dotted lines denote the boundary of the periphery of the organoids. x,y,z denote the angle of rotation. The scale bar is 500μm. Bottom panel shows a zoom-out and zoom-in view of the organoids stained with SOX2, CRYAB, GFAP. DAA denotes the days after aggregation. The scale bar is 100μm. **F**, schematics of Mon-Ch and Het-Ch mouse chimeroids. **G,** Whole-mount immunofluorescence (representative images) of SATB2 and CRYAB, 1 and 2.5 day after chimeroid aggregation. Single *z-*stack is shown. DAA is days after aggregation. N=3 organoids per condition and timepoint. The scale bar is 100μm. **H,** Quantification of SATB2 signal. Bar plot showing the proportions of SATB2-positive nuclei per ex vivo mouse organoid, grouped by three reseeding conditions: Mon-Ch (old), Mon-Ch (young), and Het-Ch from cultures 1 days (D1AM; for each group: n = 3 replicate organoids × 2 batches) and 2.5 days (D2.5AM; for each group: n = 3 replicate organoids × 1 batch) after mixing, respectively. Bars represent the mean values with error bars indicating the standard error of the mean. Jittered dots, representing the quantification of each sample, are shaped by batches. The significance of the condition effect on proportional differences across three groups at D1AM was evaluated by a likelihood ratio test comparing binomial generalized linear models with or without condition as a fixed effect while adjusting for batches (ANOVA *P*-value < 2.2 ×10^-16^). Two-sided Tukey-adjusted *P*-values for pairwise comparisons between groups were represented by asterisks.

